# A meta-analysis of effects of seaweed and other bromoform containing feed ingredients on methane production, yield, and intensity in cattle

**DOI:** 10.1101/2025.05.17.654669

**Authors:** Ermias Kebreab, Eleanor May Pressman, John-Fredy Ramirez-Agudelo, André Bannink, Sanne van Gastelen, Jan Dijkstra

## Abstract

Methane (CH_4_) emissions from ruminants contribute significantly to global GHG emissions. Bromoform (CHBr_3_)-containing feed ingredients, such as *Asparagopsis* seaweed, have emerged as promising tools to reduce enteric CH_4_ emissions. This meta-analysis quantitatively assessed the effects of CHBr_3_-containing seaweeds and synthetic CHBr_3_-based additives on CH_4_ production (g/day), CH_4_ yield (g/kg DMI), and DMI, as well as CH_4_ intensity (g/kg product) and production (milk or daily gain in dairy and beef cattle, respectively). Data were collected from 14 studies, with 39 treatment mean differences for CH_4_ production, comprising both beef and dairy cattle fed CHBr_3_-containing ingredients, while accounting for dose, DMI, and diet composition. The random-effects model’s estimates revealed that at an average CHBr_3_ dose of ≈28.3 mg/kg DM, CH_4_ production was reduced by 47.3%, CH_4_ yield by 43.3%, and CH_4_ intensity by 39.0%, with increases in CHBr_3_ dose resulting in larger efficacy in mitigating CH_4_ emissions. The efficacy of CHBr_3_ was influenced by cattle type, with greater mitigation effects in beef cattle than dairy catle, and by dietary composition, with greater reductions observed in diets higher in starch, whereas higher NDF levels attenuate its effect. At the average CHBr_3_ dose, estimated DMI was significantly reduced by 6.45% and 3.26% in dairy and beef cattle, respectively. The significant reduction in DMI did translate into a significant effect on milk yield (-4.60%) at average CHBr_3_ dose. Carrier type (oil vs. biomass), measurement technique and cattle type influenced the results. Oil carrier potentially leading to more pronounced reductions, particularly for CH_4_ intensity, respiration chambers yielded significantly greater CH_4_ reductions compared to other methods, and beef cattle showed stronger mitigation effects than dairy cattle. This study highlights the CH_4_ mitigating potential of CHBr_3_-containing feed ingredients, providing predictive models to optimize CH_4_ reduction strategies under diverse production conditions. Future research should address long-term effects, dietary optimization, and practical implementation.

## INTRODUCTION

Methane (**CH_4_**) is a potent GHG, with a global warming potential 27 or 81 times greater than carbon dioxide (**CO_2_**) over a 100-year or 20-year period, respectively (IPCC, 2021). The livestock sector, particularly large ruminant production, is a major source of anthropogenic enteric CH_4_ emission, contributing substantially to global GHG emissions (Gerber et al., 2013). Addressing enteric CH_4_ emission from cattle is critical for mitigating climate change and meeting global sustainability goals (Smith et al., 2014).

One promising approach to reduce enteric CH_4_ emission is the use of feed ingredients such as CHBr_3_-containing seaweed into cattle diets (Honan et al., 2022). Bromoform (**CHBr_3_**) is a naturally occurring compound in certain types of seaweed, such as *Asparagopsis spp.*, which has been shown to considerably inhibit enteric CH_4_ emissions during rumen fermentation (Kinley et al., 2020). The key metalloenzymes are blocked by CHBr_3_ via the Wolfe cycle (Glasson et al., 2022), inhibiting the methyl transfer of the cobamide-dependent enzyme, which is needed to catalyze the final step in methanogenesis (Fouts et al., 2022; Glasson et al., 2022). There are, however, strict safety measures for products fed to cattle that enter the human food supply, and not all CHBr_3_-containing seaweeds are FDA-approved, and in Europe CHBr_3_ extracts or otherwise are feed additives and require EFSA approval (Tricarico et al., 2025). Stabilized CHBr_3_-based additives have also been developed to achieve similar effects (e.g., Colin et al., 2024). The effectiveness of CHBr_3_-containing additives in reducing enteric CH_4_ emission has been demonstrated in several studies, with reported reductions ranging from no significant reduction to over 80% (Roque et al., 2019; Kinley et al., 2020).

However, the effectiveness of CHBr_3_-containing seaweeds and CHBr_3_-based additives can vary widely due to factors such as the dose, cattle type, diet composition, and the methods used to measure CH_4_ emission (Roque et al., 2021). Therefore, developing robust predictive equations to quantify and elucidate the factors contributing to the variation in the effects of feed additives, including CHBr_3_-containing feed ingredients, on CH_4_ emissions is essential. These equations can optimize the use of CHBr_3_-containing feed ingredients in various production systems and inform policy and management decisions to enhance sustainability in the livestock sector (Dijkstra et al., 2025).

Previous studies have attempted to quantify the effects of CH_4_-reducing additives through meta-analyses of in vivo and in vitro data (e.g., Appuhamy et al., 2013; Dijkstra et al., 2018; Martins et al., 2024). Lean et al. (2021) and Orzuna-Orzuna (2024) conducted a meta-analysis on the effects of dietary seaweed on cattle performance and CH_4_ yield, which included all types of seaweed. However, our analysis is novel in its specific focus on CHBr_3_-containing seaweeds, providing a more targeted investigation compared to the broader scope of Lean et al. (2021) and Orzuna-Orzuna et al. (2024). Although multiple compounds in *Asparagopsis* spp. may act synergistically to inhibit methanogenesis, CHBr_3_ is the most abundant natural product in *Asparagopsis* spp. (Machado et al., 2016) and the anti-methanogenic properties of red seaweed are often attributed to CHBr_3_. Variability in CHBr_3_ efficacy across studies may be due to variation in the CHBr_3_ content of seaweeds, as well as potential instability of CHBr_3_ in seaweed (e.g., Stefenoni et al., 2021; Angellotti et al., 2025). Thus, focusing our analysis on the effect of CHBr_3_ (not red seaweed per se) allows greater standardization of the impact of CHBr_3_-containing seaweed supplementation. Most studies employ frequentist statistical methods, which, although valuable, have limitations in fully capturing the complexity and uncertainty inherent in the data, particularly when dealing with limited or noisy datasets. In contrast, Bayesian methods incorporate prior knowledge and thus potentially offering a more comprehensive quantification of uncertainty (Sidebotham et al., 2023) and hence may complements these frequentist methods.

The objective of this study was to develop predictive equations that quantify the effect of CHBr_3_-containing seaweed and other CHBr_3_-based additives on CH_4_ production (g/d), yield (g/kg DM) and intensity (g/kg milk, ECM, ADG) in dairy and beef cattle. We hypothesized that supplementing CHBr_3_-containing seaweed as well as CHBr_3_-based additives reduce enteric CH_4_ production, yield, and intensity in beef and dairy cattle, and that the level of CH_4_ reduction depends on CHBr_3_ dose and diet composition. Both frequentist and Bayesian meta-analysis approaches were employed to ensure the robustness of the present study and to provide a comprehensive quantification of the uncertainty associated with the estimates of CH_4_ reduction achieved.

## MATERIALS AND METHODS

### Data Sources

A priori inclusion criteria for the studies included in the meta-analysis were defined as follows: studies must have been conducted in vivo in either dairy or beef cattle fed either CHBr_3_-containing seaweed (e.g., *Asparagopsis spp*.), a seaweed-based product, or a CHBr_3_-containing additive for the purpose of CH_4_ inhibition. The targeted and/or fed CHBr_3_ dose (mg CHBr_3_/kg DMI) must be either reported or calculable from the provided information. The CHBr_3_ products must have been fed during the same experimental timeframe as enteric CH_4_ emissions were measured and not directly introduced in the rumen. The study must have included a control group that did not receive the CHBr_3_-based treatment. Enteric CH_4_ emissions must have been measured (i.e., emission rate quantitatively assessed), not estimated through mathematical modeling, meta-analysis, or lifecycle analysis (LCA) or alike. Measurements of individual DMI must be provided. The basal diet’s chemical composition (% dietary DM) of starch, crude protein (CP), neutral detergent fiber (NDF), and crude fat (FAT) must either be reported or calculable from the basal diet’s ingredient composition. Studies must be original research articles published or under review in peer-reviewed journals (e.g., not theses, meeting abstracts, literature reviews, meta-analyses, or life cycle assessments) and written in English.

Literature searches were conducted on 1/6/2025 using Web of Science, Scopus, and CAB Abstracts databases. Search terms included combinations of (“Asparagopsis” OR “seaweed” OR “red seaweed” OR “macroalgae” OR “algae” OR “bromoform” OR “CHBr3”) AND (“dairy cattle” OR “dairy cow” OR “beef cattle” OR “steer” OR “cow” OR “cattle”) AND (“enteric methane” OR “methane” OR “CH4”). Filters were applied to include only peer-reviewed journal articles. One co-author screened abstracts and full articles as needed to assess whether the inclusion criteria were met.

The literature searches in the Web of Science, Scopus and CAB Abstracts databases yielded a total of 218, 143 and 78 entries, respectively. After removing duplicates, the remaining entries (268) underwent further screening based on the established inclusion criteria. The majority of entries (256) were excluded for the reason that studies: did not measure enteric CH_4_ emission directly; CHBr_3_ concentrations of the seaweed or CHBr_3_-based additive fed was not reported; involved in vitro experiments, bioreactor setups or co-digestion processes instead of in vivo studies; consisted of review articles, modelling studies, or meta-analyses; investigated seaweed species or other feed additives that did not contain CHBr_3_; were conducted in species other than cattle; did not involve actual feeding trials, such as surveys of farmers’ or consumers’ perceptions. A total of 12 entries met the inclusion criteria (Altman et al., 2024; Alvarez-Hess et al., 2023, 2024; Colin et al., 2024; Cowley et al., 2024; Eikanger et al., 2024; George et al., 2024; Kinley et al., 2020; Meo-Filho et al., 2024; Roque et al., 2019, 2021; Williams et al., 2024). Krizsan et al. (2023) and Stefenoni et al. (2021) studies were excluded because the CHBr_3_ concentrations in the seaweed or in the CHBr_3_-based additive fed were not reported. Kinley et al. (2024) study was excluded because SEM of CH_4_ production was not available. Although the CHBr_3_ concentration of the additive fed was not given by Colin et al. (2024), it was given in associated thesis (Colin, 2023). Hence, Colin et al. (2024) was included and CHBr_3_ concentrations from the thesis were used, but all other data were from the published article. Two manuscripts in press were verified to meet inclusion criteria and included (Angellotti et al., 2025; Kelly et al., 2025), leaving a total of 14 papers in the final dataset. The process of identifying, screening, and including studies is presented in a PRISMA (Preferred Reporting Items for Systematic Reviews and Meta-analysis; Page et al., 2021) flow diagram (Figure 1).

**Figure 1.**
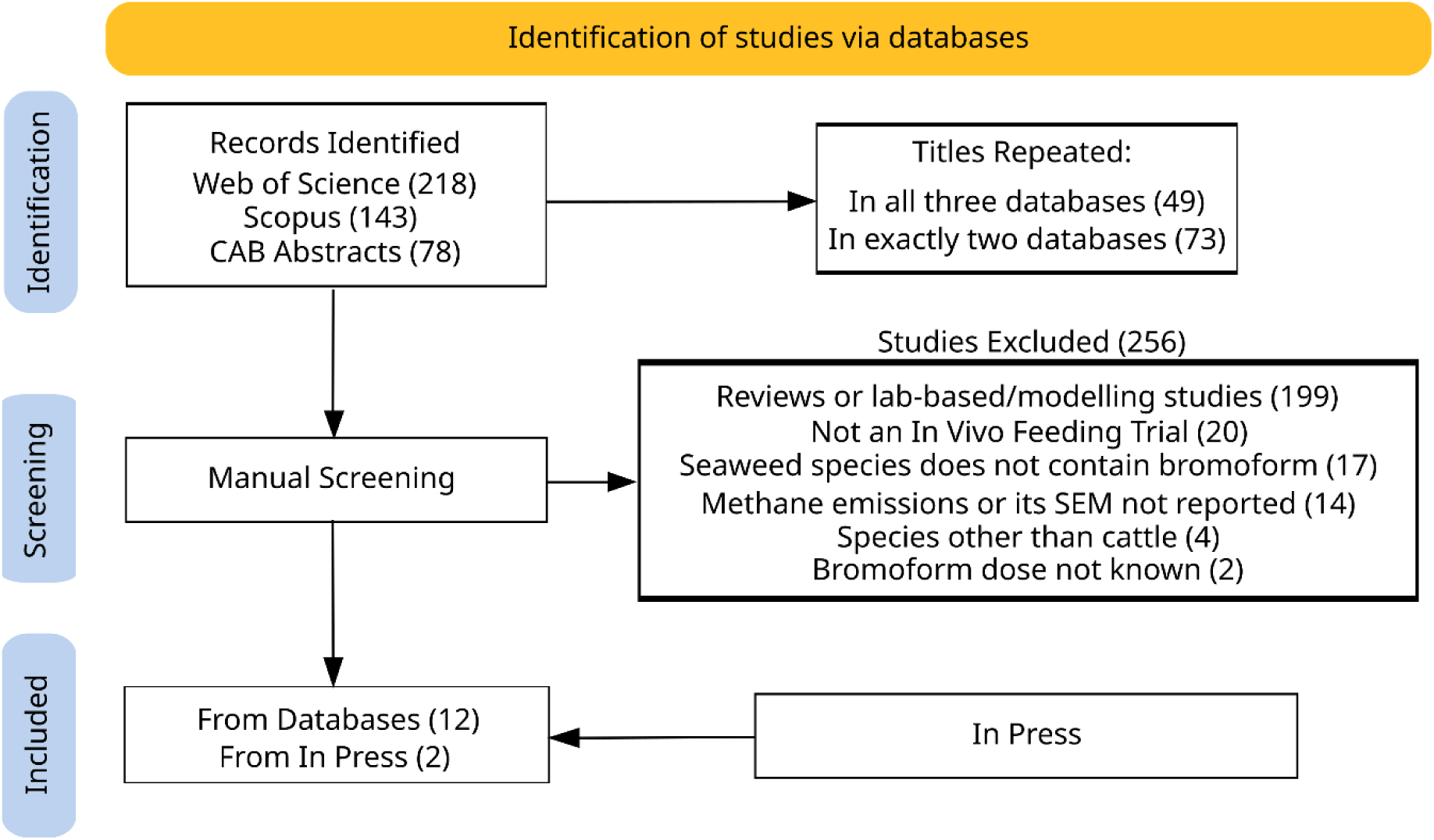
Study selection process. Literature searches were conducted using Web of Science, Scopus, and CAB Abstracts databases. Search terms included combinations of (“Asparagopsis” OR “seaweed” OR “red seaweed” OR “macroalga” OR “alga” OR “bromoform” OR “CHBr3”) AND (“dairy cattle” OR “dairy cow” OR “beef cattle” OR “steer” OR “cow” OR “cattle”) AND (“enteric methane” OR “methane” OR “CH4”).

Data were manually collected for all included studies by one co-author using an Excel database. Data collected for all studies where available were: cattle type (beef, dairy); CH_4_ production (g/day), CH_4_ yield (g/kg DM), CH_4_ intensity (g/kg milk, ECM, ADG), and their respective standard errors; average initial body weight (kg); milk and ECM yield (kg/day) or average daily gain (ADG, kg/day) and their respective standard errors; the CH_4_ measurement technique (i.e., Greenfeed, SF_6_, or respiration chamber); CHBr_3_ carrier (e.g., seaweed biomass, seaweed-steeped carrier oil, etc.). Dry matter intake and ADG (kg/d) for each treatment group and standard error were directly collected from reports of all studies except Meo-Filho et al. (2024), as described below. Bromoform intake (g/day) and CHBr_3_ dose (mg/kg diet DM) were collected or calculated for all studies as described below. Basal diet content of starch, CP, NDF, and FAT (% DM) were collected or calculated as described below for all studies.

Numerical data tables were not provided in 2 articles. The online Plot Digitizer tool (PlotDigitizer, 2024) was used to extract CH_4_ production and yield data from Figure 1 and CH_4_ intensity (g/kg milk yield) from Figure 2 in Roque et al. (2019). Personal communication with R. Kinley clarified that the error bars in figures in Roque et al. (2019) are SE (not SD, as noted in the former), so SEM for these variables was extracted by subtracting the bottom from top error bar in each column in the figures and dividing by 2. For Kinley et al. (2020), numerical data and SEM for CH_4_ production and yield, DMI (intake during CH_4_ measurement in chambers), and ADG (over treatment period) data were obtained through personal communication with the authors; SEM for CH_4_ intensity (g/kg ADG) were not available. Only predicted DMI was available for Meo-Filho et al. (2024) because of the study’s setting (grazing beef cattle). Predicted DMI and ADG (kg/day) used in CH_4_ yield and intensity calculations, and their respective SE, were obtained through personal communication with the authors. For George et al. (2024), overall trial CH_4_ emission and production data (1-200 d) were used and for Cowley et al. (2024), overall trial DMI was used. Where SEM was reported as a range in the latter study, the average of the range extremes was recorded. For Altman et al. (2024), only the data from the gas collection period in Experiment 1 were used. Experiment 2 quantified emissions reductions in steers that were previously, but not currently, supplemented with a CHBr_3_-containing feed additive, so data from this experiment were excluded. Gas emissions and SE given in L/d were converted to g/d using the ideal gas molar volume at standard temperature and pressure (**STP**; 22.4 L/mol) and gas molar mass. For gas emissions under the limit of detection in Altman et al. (2024), the limit of detection divided by the square root of 2 was used (Hornung and Reed, 1990). Production data were not available from Altman et al. (2024).

**Figure 2.**
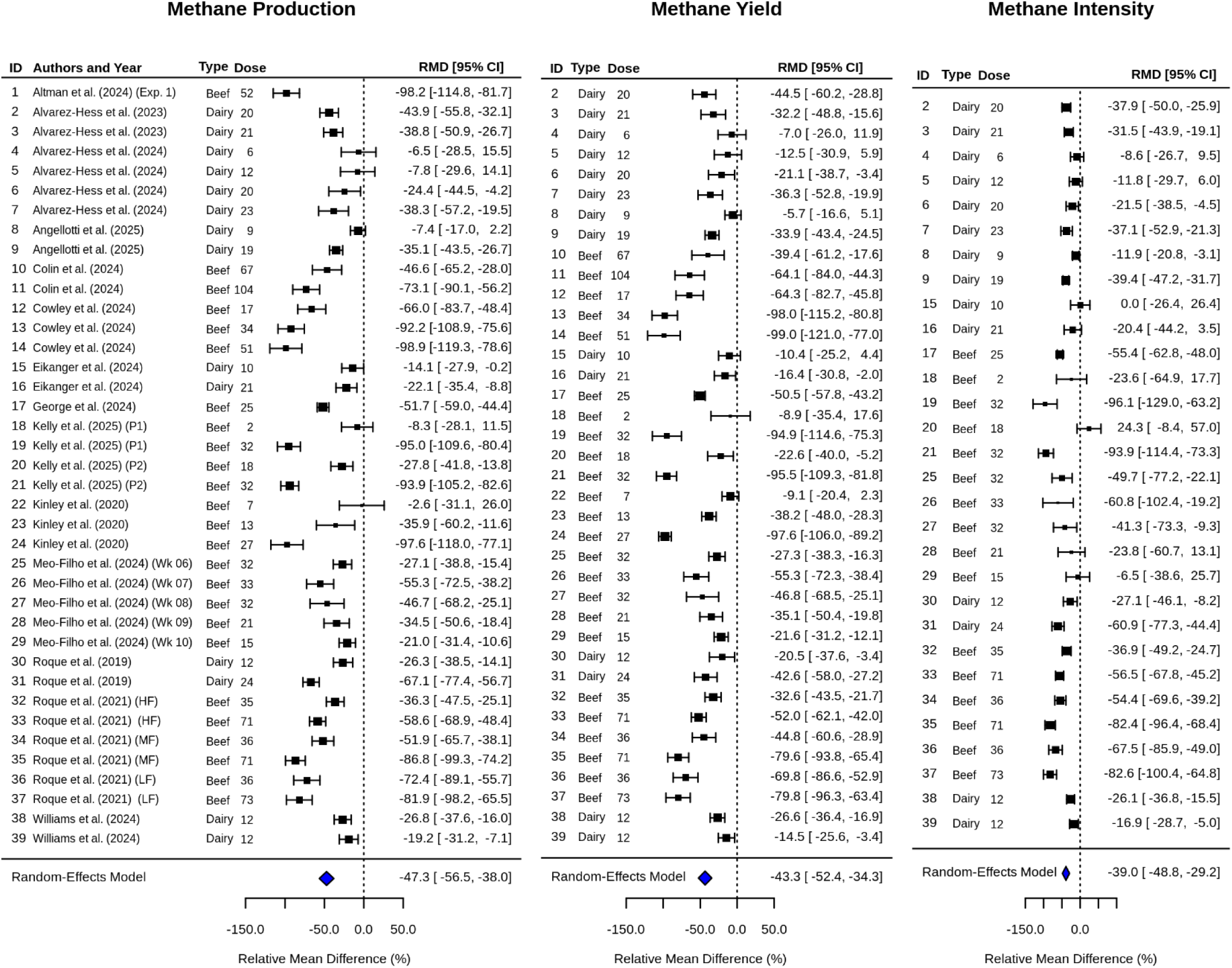
Forest plot showing bromoform (CHBr_3_) dose (mg/kg DM) and relative mean difference values (RMD is calculated as (treatment mean − control treatment mean) / control treatment mean × 100) in CH_4_ production (g/d), CH_4_ yield (g/kg DMI) and CH_4_ intensity (g/kg milk, ECM or ADG) for beef and dairy cattle studies. P1 and P2 = experimental periods; Wk = experimental week; HF = high forage diet; LF = low forage diet; MF = medium forage diet. The black squares represent the power of the study (i.e., greater sample sizes and smaller confidence intervals are indicated by a larger box). The summary values at the bottom of the Forest Plot (the diamond, showing -47.3%, -43.3%, -39.0%) represent the pooled estimate from the Random-Effects Meta-Analysis model.

Doses of CHBr_3_ in Eikanger et al. (2024) and Roque et al. (2019, 2021), where seaweed was fed on an OM basis, were calculated by first multiplying the reported seaweed application rate, provided on an OM basis (g seaweed OM/g diet OM), by the OM content of the basal diet DM. This product was then multiplied by the reported CHBr_3_ concentration of the seaweed on a DM basis, and then by the OM content of the seaweed DM, to give (mg CHBr_3_ /kg diet DM). Although seaweed was also fed on an OM basis in Angellotti et al. (2025), seaweed OM content was not reported, so CHBr_3_ doses reported in the article’s discussion were used as follows: the average of 20.7 and 16.8 mg CHBr_3_/kg DMI was used for the high group dose, and half of this was used for the low group dose.

Total CHBr_3_ ingested (g/d) was determined by multiplying the CHBr_3_ dose by the DMI for each CHBr_3_-treated group. In Kinley et al. (2020), the CHBr_3_ dose was calculated by multiplying the daily as-fed TMR intake for each group by the *Asparagopsis taxiformis* inclusion rate in the as-fed ration. This result was then multiplied by the CHBr_3_ content of the as-fed *A. taxiformis* to determine the daily CHBr_3_ intake (g/d). Finally, the daily CHBr_3_ intake was divided by each treatment group’s daily DMI to express CHBr_3_ dose as mg CHBr_3_/kg DM. In Alvarez-Hess et al. (2024, 2023) and Williams et al. (2024), CHBr_3_ intake was calculated by multiplying the Asp-Oil intake by its CHBr_3_ concentration. The CHBr_3_ dose was then calculated by dividing the CHBr_3_ intake by the DMI. The online Plot Digitizer tool was also used to extract per-week CHBr_3_ intake (mg/day) from Figure 1 in Meo Filho et al. (2024). Bromoform dose was calculated by dividing CHBr_3_ intake by DMI.

In studies where grain was delivered via a concentrate station (Alvarez-Hess et al., 2023, 2024; Williams et al., 2024), the composition of the overall ingested diet was calculated by weighting each chemical component (starch, CP, NDF, and FAT as % DM) of the forage and concentrate portions according to their respective contribution to the treatment group’s overall DMI. Because dietary chemical composition was not provided by Altman et al. (2024), starch, CP, NDF, and FAT content of the overall experimental diet was calculated by multiplying the starch, CP, NDF, and FAT content of each ingredient given by NASEM (2021) by the proportion of that ingredient in the diet and then summing for all ingredients. Dietary starch content was calculated in the same manner for George et al. (2024) and Kelly et al. (2025). For George et al. (2024), information on nutrient composition was from the analyzed ration. Starch content for Meo-Filho et al. (2024) was assumed to be 2%, based on the starch content of the pellet ingredients, as the steers’ diet consisted of grass, which contributes negligible starch to the overall intake. The CP, NDF, and starch content of the diet in Angellotti et al. (2025) was determined by dividing the daily intake of each component by the total DMI for each treatment group. Because FAT intake was not reported, the FAT content of the basal diet was calculated by multiplying the FAT content of each ingredient given by (NASEM, 2021) by the proportion of that ingredient in the diet and then summing for all ingredients. Given they were fed a finishing diet, the nonlactating, nonpregnant Jersey cattle in Colin et al. (2024) were categorized as beef. Production data were not available from this study.

### Effect size calculation

The data were analyzed using methods similar to those of Kebreab et al. (2023), focusing on a frequentist meta-analysis of CH_4_ emission. This analysis centered on 2 fundamental calculations: the effect size and its variance. We defined our custom effect size as the relative mean difference (**RMD**) in CH_4_ production (g/d), CH_4_ yield (g/kg DMI), CH_4_ intensity (g/kg milk, ECM, ADG), DMI (kg/d), and product output (kg/d milk, ECM, ADG), expressed as the percentage change from the control to the treatment group. This approach allowed for comparisons across studies with varying baseline values. Importantly, whether CH_4_ intensity is measured per milk output, per ECM, or per ADG, the analysis outcome remains consistent. This consistency is because the RMD framework inherently accounts for baseline variations. As a result, it ensures comparability between dairy and beef systems by standardizing proportional changes between treatment and control groups. For example, Roque et al. (2019) report CH_4_ yield per milk output rather than per ECM. Nevertheless, the RMD framework guarantees consistency in effect size calculations across different measurement units. The RMDs were calculated using the equation:

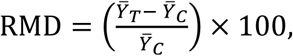

where 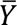 = mean for the treatment (T) and control (C) groups. Since the calculation of RMD is essentially a ratio of 2 random variables, the variance calculation had to account for the variability in both the numerator and the denominator. We applied the Delta Method (e.g., Deng et al., 2018), using the same underlying statistical principle employed by Veneman et al. (2016) and Purba and Sangsawad (2025) for the variance of the response ratio, to ensure the variability in the treatment and control groups was accurately incorporated into the RMD uncertainty. We assumed that the groups in the RMD were independent, with zero covariance between their means. The variance was calculated using the following equation:

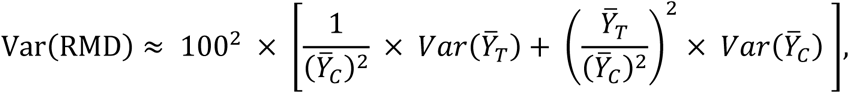

where 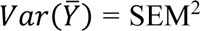 for the treatment (T) and control (C) group means, assuming that the reported SEM represents the variability within each group (Ramirez-Agudelo and Kebreab, 2024). The equation takes the uncertainty in the treatment group 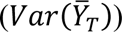 and scales it by how much the control group influences the RMD 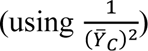. It also takes the uncertainty in the control group 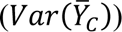 and scales it by how much the interaction of the treatment and control values affects the RMD 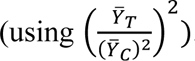. These 2 contributions are then added to combine the total uncertainty. Finally, the result of uncertainty of RMD (Var(RMD)) is obtained by multiplying by 100^2^because the RMD is expressed as a percentage. The treatment group’s influence is simpler because it only affects the numerator in the RMD calculation, while the control group’s influence is more complex because it affects both the numerator and denominator. The sampling variances are crucial for weighting the effect sizes in the meta-analysis. Studies with smaller variances (more precise estimates) are given more weight in the analysis than studies with larger variances (Viechtbauer, 2010).

### Statistical analysis

Both frequentist and Bayesian approaches were employed to meta-analyze the database and quantify the uncertainty associated with the effect of covariates on the reduction of CH_4_ emission. We evaluated several categories: cattle type (included as 2 binary category: Dairy was coded as 1 if the animal was dairy cattle and as 0 if otherwise, and Beef was coded as 1 if the animal was beef cattle and as 0 if otherwise), CHBr_3_ dose (mg/kg DM), quadratic CHBr_3_ dose ((CHBr_3_ dose)^2^), DMI (kg/d), starch (% of DM), CP (% of DM), NDF (% of DM), and FAT (% of DM). In models that included starch, data from Roque et al. (2019) were excluded because the starch proportion was not reported and dietary data were not available to calculate it. Covariates, except cattle type, were centered within each cattle type by subtracting their mean values to reduce multicollinearity, specifically between CHBr_3_ dose and its squared term, and to improve model’s results interpretability. Before centering, natural log transformation was applied to CHBr_3_ to reduce its skewness (Supplemental Figure S1). This transformation normalizes the distribution, particularly for beef, which exhibited significant skewness (Shapiro-Wilk *P*-value = 0.031 before transformation). Equations derived from our analysis assume that CHBr_3_ doses are first natural log-transformed. Using untransformed data in these equations would yield incorrect results, as the relationships were defined on the log scale.

Biological systems often exhibit diminishing returns at higher treatment doses, where the effect plateaus or even reverses due to physiological saturation or adverse impacts (e.g., Calabrese et al., 2003). To capture this behavior, we included the (natural log of CHBr_3_ dose)^2^ term as covariate. We assumed that both the linear and the quadratic terms are important because they represent different biological processes: the linear term captures the direct inhibitory effect of CHBr_3_, whereas the quadratic term reflects saturable effect of CHBr_3._ Including both ensures the model is both biologically realistic and mathematically robust.

In our frequentist meta-analysis, we explored all possible combinations of up to 5 covariates. These models consistently included cattle type (i.e., the 2 binary covariates: Dairy and Beef) and natural log of CHBr_3_ dose as primary predictors, along with 1, 2 or 3 additional covariates as fixed effects. The models were structured to include an intercept-free formulation via the mods argument in the ‘rma’ function (i.e., mods = ∼ 0 + Dairy + Beef + CHBr_3_ + additional covariates), using Maximum Likelihood (ML) estimation. To avoid multicollinearity, among covariates in the model, Pearson correlation coefficients were calculated, and any model containing covariate pairs with an absolute correlation coefficient greater than 0.50 was excluded. Each candidate model was fitted using mixed-effects meta-regression analysis with the ‘rma’ function from the ‘metafor’ package (Viechtbauer, 2010) in R (R Core Team, 2021). Many studies in our database included multiple treatment groups compared to a common control group, likely resulting in correlations between observations within each study. To address this within-study correlation, we applied Robust Variance Estimation (**RVE**), clustered by study, using the ‘robust’ function.

Models were evaluated based on Akaike and Bayesian information criteria (**AIC**, **BIC**), concordance correlation coefficient (**CCC**) between reported and predicted RMDs, and the root mean square error (**RMSE**) of leave-one-out cross-validation (**LOOCV**; Allen, 1974), ensuring statistical significance (*P* ≤ 0.05) for all covariates. During model fitting, Cook’s distances (Cook, 1977), calculated with the ‘cooks.distance’ function, were used to reveal outliers which influenced the regression (i.e., how much the estimated regression coefficients change when a particular observation is removed from the dataset). A common rule of thumb is to consider observations as influential if their Cook’s distance values exceed 4/N, where N is the total number of observations (e.g., Altman and Krzywinski, 2016). However, applying this to our database resulted in the removal of an excessive quantity of data points (between 25 to 55%, depending on the model structure), which would lead to a considerable loss of information and reduce the robustness of the analysis. To address this issue, we adopted a data-driven approach by defining the threshold as 3 times the mean Cook’s distance. This method provided a more balanced criterion for identifying influential data points whilst preserving the integrity of the database, and is used in the main document of this study. Cook’s distance-based outlier identification was performed separately for each candidate model. Because a data point’s influence depends on the model’s covariate set, the specific outliers identified varied slightly across models.

To evaluate the sensitivity of the results to outlier elimination, we also implemented an approach using the CCC metric as an alternative to the Cook’s distance approach. Unlike Cook’s distance, which identifies outliers based on their influence on regression coefficients, the CCC-based method targets observations with the largest residuals (i.e., discrepancies between predicted and actual values of the response variable, RMD). The algorithm iteratively removes the most extreme residual (in absolute value), refits the model, and recalculates predictions until the number of removed data points matches the count identified by Cook’s distance in the respective model. Critically, the CCC method does not necessarily remove the same data points as Cook’s distance, as it prioritizes prediction accuracy rather than coefficient stability. The results obtained with the CC metric in the outlier elimination method are shown in the supplementary document of this study.

To estimate how well each model might generalize and predict outcomes on new, unseen data, we calculated the RMSE derived from LOOCV. In the RMSE-LOOCV process, after outliers elimination, 1 observation was omitted at a time, the model was refitted using the remaining data, and the omitted observation was predicted. The RMSE was calculated for each prediction, and the average RMSE across all iterations. Additionally, the amount of residual heterogeneity (*τ^2^*) and its proportion (*I^2^*) were extracted from the summary of the models. The degree of heterogeneity measured by the *I^2^* statistic can be classified as follows: no heterogeneity (0% < *I^2^* ≤ 25%), low heterogeneity (25% < *I^2^*≤ 50%), moderate heterogeneity (50% < *I^2^* ≤ 75%), and high heterogeneity (*I^2^* > 75%) (Borenstein et al., 2017).

After removing outliers detected with the Cook’s distance method, we constructed funnel plots to visually assess the presence of publication bias (Appuhamy et al., 2013). Our analyses involved 2 types of meta-analytic models, both fitted with the ‘rma’ function. First, we estimated a random-effects model to obtain an overall pooled effect size while accounting for between-study heterogeneity. We specified the observed effect sizes (RMD) and their variances (i.e., res_random <-rma(yi = RMD, vi = var(RMD), data = data, method = “ML”)). Next, we fitted a mixed-effects meta-regression model using the same function but including cattle type (i.e., the 2 binary covariates: Dairy and Beef), log-centered CHBr_3_ dose and centered starch as moderators (i.e., res_mixed <-rma(yi = RMD, vi = var(RMD), mods = ∼ 0 + Dairy + Beef + log-centered CHBr_3_ + centered starch, data = data, method = “ML”)). Egger’s regression test was performed using the ‘regtest’ function on both type of models.

To determine the effect of CHBr_3_ supplementation on RMD of DMI and products (milk, ECM, or ADG), we employed a similar meta-analytic mixed model approach as used for the CH_4_ outcomes, utilizing the ‘rma’ function. The used model included the 2 binary covariates (Dairy and Beef) and the CHBr_3_ dose term, centered within each cattle type. Key differences for the DMI and products analysis were that the CHBr_3_ dose was used on its original scale (mg/kg DM) without log transformation, and only the model containing the linear CHBr_3_ term was evaluated. The relationship between CHBr_3_ dose and production outcomes was assumed to be linear within the studied dose range, unlike CH_4_ outcomes which showed a nonlinear response. Pairwise differences in RMD for CH_4_ production, yield, and intensity between carrier methods (biomass vs. oil), techniques (respiration chamber vs. Greenfeed vs. SF_6_), and cattle type (dairy vs. beef) were estimated using a mixed-effects meta-regression model. To get a holistic view of the combined influence of factors, a single comprehensive model was fitted for each CH_4_ outocome including carrier, technique, cattle type, and log-centered CHBr_3_ dose in the model (i.e., mods <-∼ Carrier + Technique + Cattle Type + log-centered CHBr_3_) using the ‘rma’ function. The model parameters were estimated using restricted maximum likelihood (REML). Cluster-robust standard errors were calculated, clustering by Study, using the ‘robust’ function (i.e., RVE). Pairwise differences between the levels of technique, carrier, and cattle type were then conducted using estimated marginal means (emmeans package; Lenth, 2022), applying Tukey’s method for multiple comparison adjustment, based on the fitted robust model. Some studies were excluded from this part of the analysis due to the limited data available on the reported types of carriers and technique. Specifically, Two papers using the headbox-type indirect calorimeter technique (Altman et al., 2024; Colin et al., 2024) were not used in this part of the analysis. Two studies using respiration chambers (Cowley et al., 2024; Kinley et al., 2020), did not report SEM of CH_4_ intensity and were therefore not included in the analysis on CH_4_ intensity for technique comparisons. Altman et al. (2024) utilized a Proprietary Kelp Blend Product as the carrier of CHBr_3_, whereas Kelly et al. (2025) employed a powder carrier in 1 treatment. Although Colin et al. (2024) did not report specific details about the carrier within the bromoform containing product (Alga 1.0) used, we classified this study in the biomass carrier group.

Finally, in our Bayesian analysis, the ‘brms’ package (Bürkner, 2017) in R was employed to perform a comprehensive Bayesian meta-analysis to quantify the uncertainty surrounding the effects of (LN(CHBr3 dose))^2^, DMI and dietary covariates on CH_4_ production, yield, and intensity. This analysis leveraged the flexibility and robustness of Bayesian modeling to provide probabilistic interpretations of parameter estimates and their associated uncertainties. The 6 covariates (LN(CHBr_3_ dose))^2^, DMI, starch, CP, NDF, and FAT) were evaluated across the CH_4_ reduction outcomes. A natural log transformation was applied to CHBr_3_ concentration (mg/kg DM) and used as co-variate, and covariates were mean-centered within each cattle type. The 6 Bayesian models were structured with the RMD as the response variable, alongside its variance, which was explicitly modeled using the ‘se’ function. Fixed effects included the 2 cattle type binary covariates (Dairy and Beef) and LN(CHBr_3_ dose), as well as one of the 6 covariates. Random effects were incorporated to explicitly models between-study heterogeneity through a random intercept term (1 | Study). The formula for the Bayesian model was structured as: RMD | se(sqrt(var (RMD)), sigma = TRUE) ∼ 0 + Dairy + Beef + LN(CHBr_3_ dose) + one of the 6 covariates + (1 | Study).

The Bayesian modeling was implemented using the ‘brm’ function with 4 Markov chains to ensure robust sampling. Each chain consisted of 10,000 iterations, including a 3,000-iteration warm-up (burn-in) period. Convergence was facilitated by setting the control parameter adapt-delta to 0.999, and a fixed random seed (i.e., 123) ensured reproducibility. Covariate-specific models results were saved as RDS files for further analysis. Key metrics, including posterior means, credible intervals, and R-hat values, were examined to assess model convergence and the reliability of the results (Bürkner, 2017).

## RESULTS AND DISCUSSION

### Methane Production, Yield, and Intensity

The meta-analysis assessed the effectiveness of CHBr_3_-containing seaweeds and CHBr_3_-based feed additives on CH_4_ emission in both dairy and beef cattle, using a database covering a range of experimental conditions, doses, DMI, and key dietary covariates. The database comprised information from 14 studies, with 39 unique comparisons between experimental and control groups. Among these, 6 studies were conducted on dairy cattle and 8 on beef cattle. Altman et al. (2024) reported only CH_4_ production, Colin et al. (2024) did not report CH_4_ intensity, and Cowley et al. (2024) and Kinley et al. (2020) lack SEM data for CH_4_ intensity. Consequently, 38 treatment comparisons were included in the analysis of CH_4_ yield, and 30 were used for CH_4_ intensity. Table 1 presents descriptive statistics of studies, the mean and SEM for CHBr_3_ dose, DMI, dietary components, CH_4_ emissions and production outcome, categorized into 3 groups: all data (encompassing both dairy and beef cattle) across the 3 CH_4_ outcomes, dairy only, and beef only. Table 2 presents descriptive statistics of studies, the mean and SEM for CHBr_3_ dose, DMI, and product output (milk, ECM, or ADG) again categorized into 3 groups: all data (encompassing both dairy and beef cattle), dairy only, and beef only. The results of the Random-Effects Meta-Analysis model for CH_4_ outcomes are presented in Figure 2. The RMD values for CH_4_ production and yield were all negative, and for CH_4_ intensity only 2 data points comparisons—Eikanger et al. (2024; low dose level) and Kelly et al. (2025, low dose level in P2)—had RMD values of 0 and +24.3%, respectively. This indicates that CHBr_3_ supplementation (i.e., in the form of seaweed or stabilized feed additives) generally demonstrated an anti-methanogenic effect.

**Table 1.**
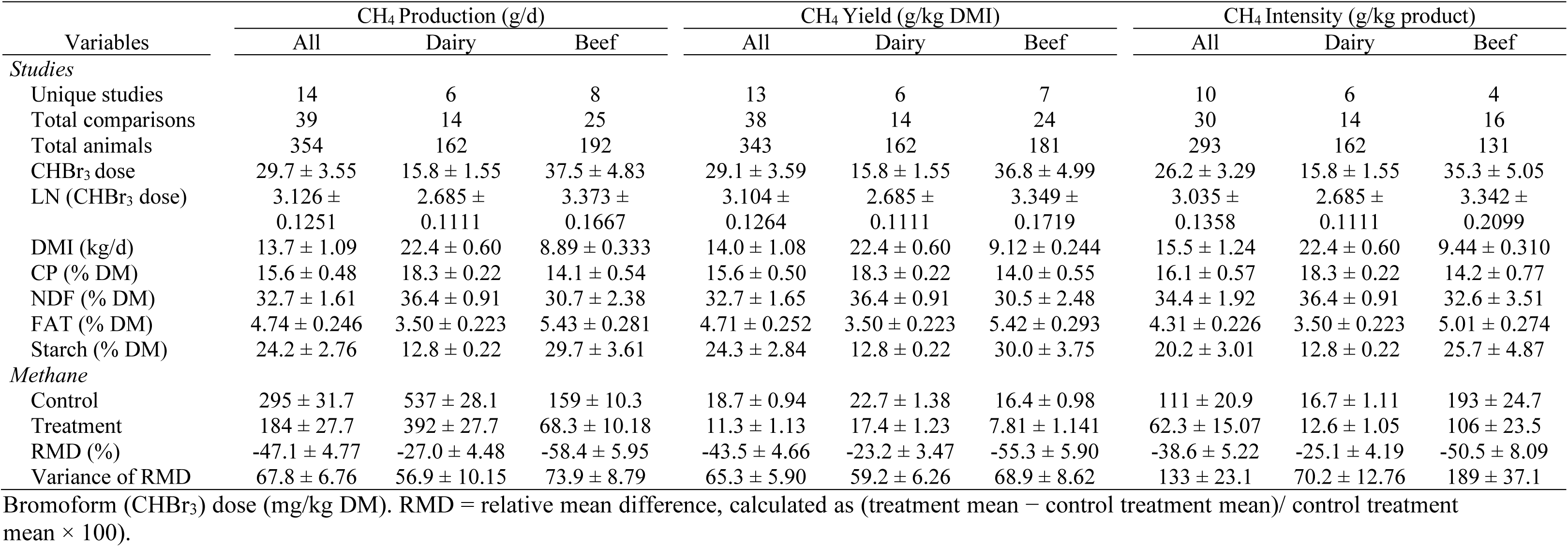
Descriptive statistics of studies and CH_4_ emission for All (encompassing both dairy and beef cattle), Dairy, and Beef across the 3 CH_4_ outcomes datasets (mean ± SEM)

**Table 2.** Descriptive statistics of studies, DMI (categorized into 3 groups: all data, dairy only, and beef only), and product output (categorized into dairy or beef) mean ± SEM)

| Variables | DMI<br>(kg/d) |  |  | Product<br>(milk, ECM, ADG kg/d) |  |
| --- | --- | --- | --- | --- | --- |
|  | All | Dairy | Beef | Dairy | Beef |
| *Studies* |  |  |  |  |  |
| Unique studies | 14 | 6 | 8 | 6 | 6 |
| Total comparisons | 39 | 14 | 25 | 14 | 22 |
| Total animals | 354 | 162 | 192 | 162 | 169 |
| CHBr<sub>3</sub> dose | 29.7 ± 3.55 | 15.8 ± 1.55 | 37.5 ± 4.83 | 15.8 ± 1.55 | 32.4 ± 4.13 |
| *DMI or Product* |  |  |  |  |  |
| Control | 14.8 ± 1.21 | 24.4 ± 0.48 | 9.36 ± 0.386 | 33.2 ± 0.90 | 1.27 ± 0.073 |
| Treatment | 13.7 ± 1.09 | 22.4 ± 0.60 | 8.89 ± 0.333 | 32.0 ± 1.04 | 1.30 ± 0.081 |
| RMD (%) | -5.54 ± 1.582 | -7.93 ± 2.630 | -4.20 ± 1.969 | -3.73 ± 1.533 | 3.16 ± 3.754 |
| Variance of RMD | 38.1 ± 6.65 | 25.9 ± 4.03 | 44.9 ± 9.94 | 27.1 ± 6.54 | 476 ± 189 |

Bromoform (CHBr<sub>3</sub>) dose (mg/kg DM). RMD = relative mean difference, calculated as (treatment mean – control treatment mean)/ control treatment mean × 100).

The detection and removal of outliers in this study were critical to ensure the accuracy and reliability of the results. By identifying and eliminating these outliers, we ensured that any lack of significant effect observed in the analysis was not due to the distortion caused by anomalous values, but rather to the absence of a true effect.

The results of this study are presented in 3 parts. The primary analysis, which uses Cook’s distance to identify and address outliers, is detailed in the main document (Tables 3, 4, and 5). To offer a more comprehensive perspective, 2 supplementary analyses are included in the supplementary material. The first supplementary analysis shows the results without any outlier removal (Supplemental Table S1 to Table S4), serving as a baseline to assess the influence of outliers on the outcomes. The second supplementary analysis (Supplemental Table S5 to Table S8) applies the CCC-based method for outlier removal, providing an alternative approach to handling outliers. Together, these supplementary analyses highlight the sensitivity of the results to different outlier treatment methods, allowing readers to evaluate the robustness and reliability of our conclusions across different methodological approaches. Cook’s distance was selected as the primary outlier diagnostic because it quantifies each observation’s influence on regression coefficients, ensuring model stability (Cook, 1977). This standard regression tool (e.g., Díaz-García and González-Farías, 2004) combines leverage and residual information to identify observations that disproportionately affect model predictions.

**Table 3.** Comparison of regression models evaluating the effect of bromoform dose (CHBr_3_; in LN (mg CHBr_3_/ kg DM) and additional dietary (% DM) or animal covariates (cattle type) on the relative mean difference in CH_4_ production (estimates ± SE)

| Models | Estimates |  |  |  |  | Model metrics |  |  |  |  |  |
| --- | --- | --- | --- | --- | --- | --- | --- | --- | --- | --- | --- |
|  | Dairy | Beef | 2nd Covariate | 3rd Covariate | 4th Covariate | Outliers ID | *I*<sup>2</sup> | AIC | BIC | CCC | RMSE (%) |
| Type + CHBr<sub>3</sub> | -25.1 ± 2.22<br>*P* < 0.001 | -57.3 ± 7.02<br>*P* < 0.001 | -25.9 ± 5.86<br>*P* = 0.001 | NA | NA | 18, 31 | 85.2 | 329 | 336 | 0.745 | 19.9 |
| Type + CHBr<sub>3</sub> + Starch | -24.8 ± 2.47<br>*P* < 0.001 | -61.2 ± 5.22<br>*P* < 0.001 | -26.6 ± 3.75<br>*P* < 0.001 | -0.764 ± 0.1525<br>*P* = 0.001 | NA | 10, 11 | 76.1 | 298 | 306 | 0.863 | 16.4 |
| Type + CHBr<sub>3</sub> + NDF | -27.5 ± 3.13<br>*P* < 0.001 | -61.9 ± 7.32<br>*P* < 0.001 | -26.9 ± 5.35<br>*P* = 0.001 | 0.839 ± 0.1934<br>*P* = 0.002 | NA | 10, 11, 25 | 80.0 | 312 | 319 | 0.833 | 17.6 |
| Type + CHBr<sub>3</sub> + (CHBr<sub>3</sub>)<sup>2</sup> + NDF | -29.9 ± 3.36<br>*P* < 0.001 | -63.5 ± 7.40<br>*P* < 0.001 | -23.7 ± 5.23<br>*P* = 0.001 | 15.7 ± 6.68<br>*P* = 0.043 | 0.848 ± 0.2764<br>*P* = 0.013 | 18 | 81.6 | 334 | 343 | 0.803 | 18.8 |

The covariates follow the same order as they are presented in the models. The first covariate corresponds to the cattle type which must be one of the two columns: Dairy or Beef. For example, in the model 'Type + CHBr<sub>3</sub> + Starch', the '2nd Covariate' column shows the estimate for the (natural log-transformed, centered) CHBr<sub>3</sub> dose, and the '3rd Covariate' column shows the estimate for (centered) Starch.
Type = cattle type (dairy or beef). Cattle type was included as 2 binary category in the models: Dairy was coded as 1 if the animal was dairy cattle and as 0 if otherwise, and Beef was coded as 1 if the animal was beef cattle and as 0 if otherwise.
All variables (except cattle type) were centered to their mean value within each cattle type presented in Table 1.
Bromoform (CHBr<sub>3</sub>) dose (mg/kg DM) was natural log-transformed before models fitting.
*I*<sup>2</sup> = proportion of total variability due to heterogeneity.
AIC, BIC = Akaike and Bayesian information criteria.
CCC = Concordance Correlation Coefficient.
RMSE = root mean square error using leave-one-out cross-validation, quantifying how far the model's predictions typically deviate from the actual observed values, in percentage points of relative mean difference (RMD).

**Table 4.**
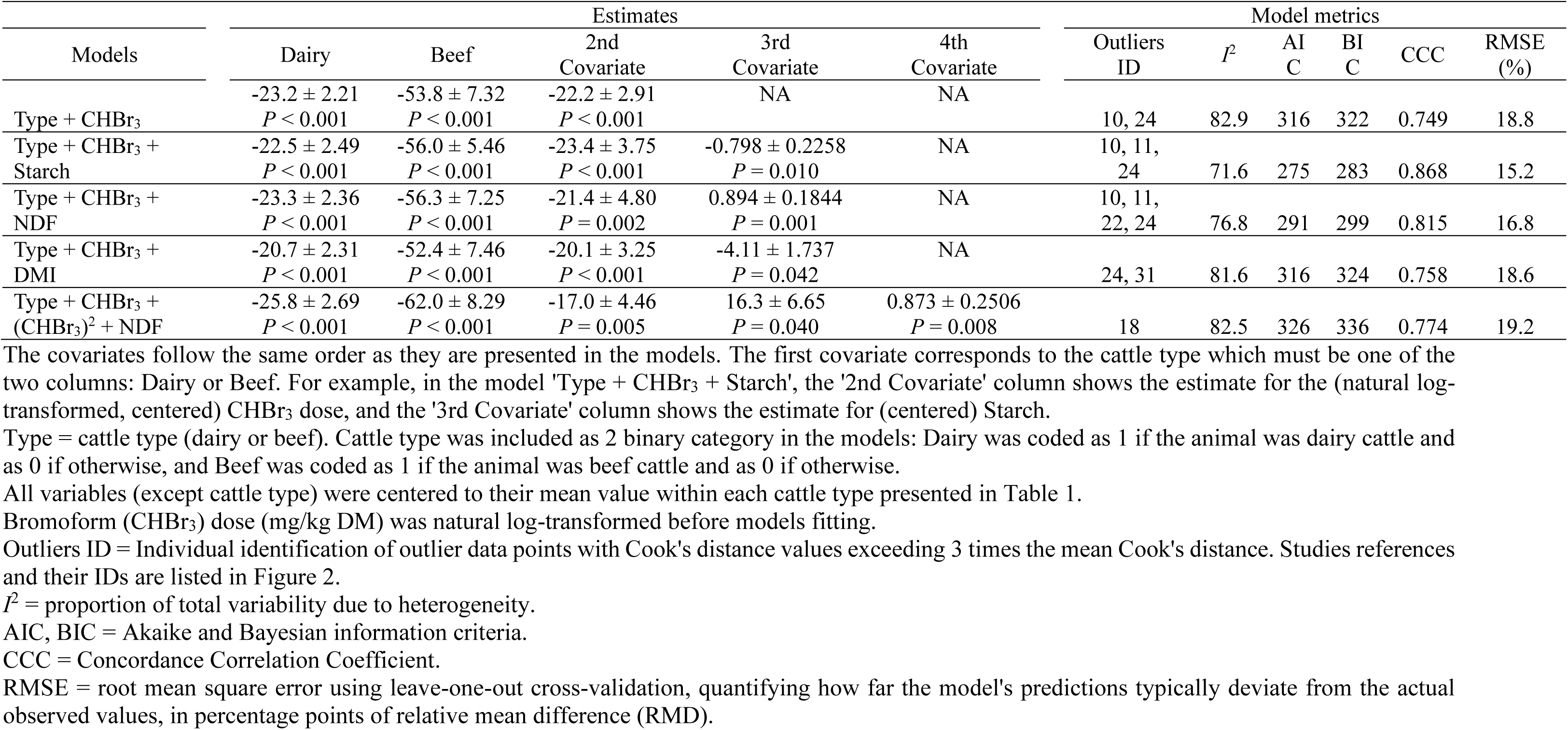
Comparison of regression models evaluating the effect of bromoform dose (CHBr3; in LN mg CHBr3/kg DM) and additional dietary (% DM) or animal covariates (cattle type) on the relative mean difference in CH_4_ yield (estimates ± SE)

Our analysis revealed distinct patterns in the significance of variables related to CH_4_ production, yield, and intensity, depending on the treatment of outliers. When outliers were not removed, only NDF was a significant covariate (*P* < 0.05) for CH_4_ production and yield, and starch exhibited a trend (*P* < 0.10). For CH_4_ intensity, NDF, starch, and (LN(CHBr_3_ dose))^2^ were significant covariates (*P* < 0.05). However, when the CCC-based method was applied to eliminate outliers, the significance of other covariates emerged (Supplemental Tables S5, S6, and S7). For CH_4_ production, FAT, starch, and NDF were significant (*P* < 0.05); for CH_4_ yield, FAT and starch were significant (*P* < 0.05), and NDF showed a trend (*P* = 0.071); and for CH_4_ intensity, starch were significant (*P* < 0.05); and LN (CHBr_3_)^2^ and NDF showed a trend (*P* < 0.10).Tables 3 to 5 present the results obtained using Cook’s distance, the reference method for outlier detection, for CH_4_ production, yield and intensity, respectively. When utilizing the estimators presented in the tables (in both the main document and the supplementary material), 2 key considerations must be addressed to ensure accurate application. First, the tables include separate columns for Dairy and Beef, which are mutually exclusive. This design requires that the appropriate column value be selected based on the cattle type under analysis. Specifically, equations intended for dairy cattle must exclusively incorporate values from the Dairy column, while those for beef cattle should use values from the Beef column. Second, the CHBr_3_ dose as reported in studies in mg/kg DM must be natural log-transformed prior to its inclusion in any of the CH_4_ emission calculations made with the equations presented here. This transformation is essential because all models were developed and validated based on this natural log scale. Adhering to these guidelines is critical for the proper interpretation and application of the reported estimators.

**Table 5.**
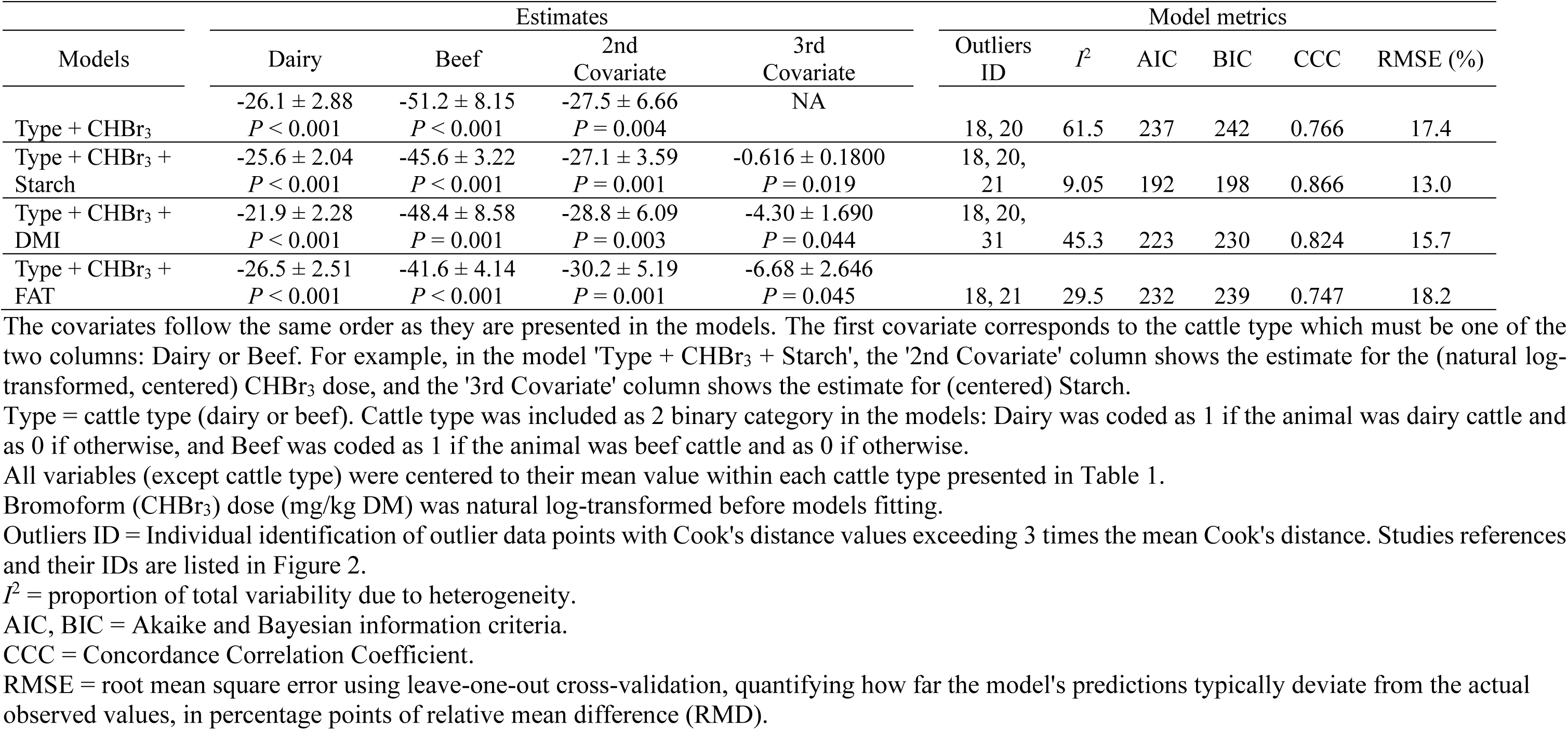
Comparison of regression models evaluating the effect of bromoform dose (CHBr3; in LN mg CHBr3/kg DM) and additional dietary (% DM) or animal covariates (cattle type) on the relative mean difference in CH_4_ intensity (estimates ± SE)

The model selection process identified key dietary characteristics influencing CHBr_3_ efficacy. In CH_4_ production, the starch-inclusive model was the most impactful, in terms of model efficiency, as evidenced by the lowest AIC (298), BIC (306), LOOCV-RMSE (16.4%), and *I^2^* values (76.1%), and highest CCC (0.863) value. RMSE is expressed as a percentage of the observed mean, representing the average magnitude of prediction errors relative to the actual values.. Similar was observed for CH_4_ yield, were the starch-inclusive model (Type + CHBr_3_ + Starch) emerged as the most impactful, with the lowest AIC (275), BIC (283), LOOCV-RMSE (15.2%), and *I^2^* values (71.6%), and highest CCC (0.868) value. In line with these results, also for CH_4_ intensity, the starch-inclusive model demonstrated the best good fit statistics with the lowest AIC (192), BIC (198), low LOOCV-RMSE (13.0%), and *I^2^* values (9.05%), and highest CCC (0.866) value. NDF was not a significant (*P* = 0.076) explanatory variable for CH_4_ intensity in any model that included this variable. Below we present the best models (i.e., lowest AIC, BIC, and RMSE-LOOCV and highest CCC values) to predict the percentage change in CH_4_ production, yield, and intensity using the within each cattle type means reported in Table 1:

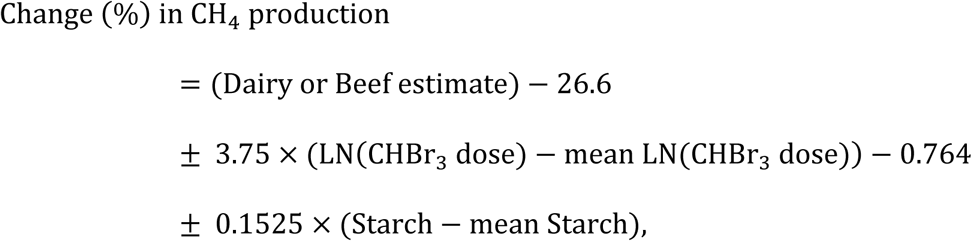

where the estimates for Dairy or Beef are -24.8 ± 2.47 or -61.2 ± 5.22, respectively; the mean LN(CHBr_3_ dose in mg/kg DM) for dairy and beef are 2.685 ± 0.1111 and 3.373 ± 0.1667, respectively; and the mean starch content (% DM) for dairy and beef are 12.8 ± 0.22 and 29.7 ± 3.61, respectively.

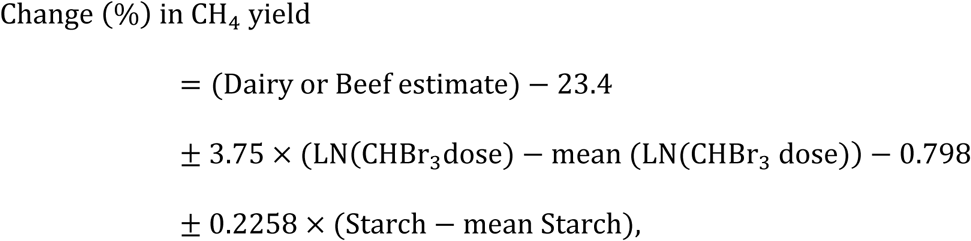

where the estimates for Dairy or Beef are -22.5 ± 2.49 or -56.0 ± 5.46, respectively; the mean LN(CHBr_3_ dose in mg/kg DM) for dairy and beef are 2.685 ± 0.1111 and 3.349 ± 0.1719, respectively; and the mean starch content (% DM) for dairy and beef are 12.8 ± 0.22 and 30.0 ± 3.75, respectively.

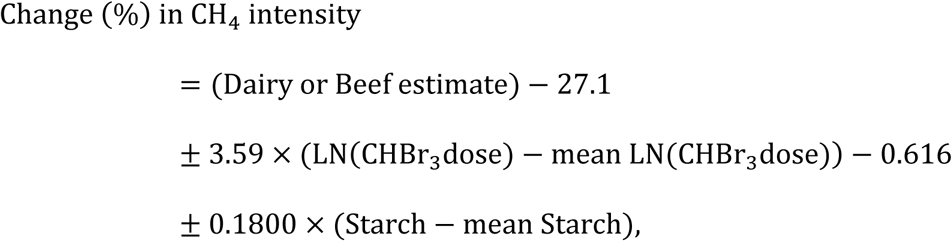

where the estimates for Dairy or Beef are -25.6 ± 2.04 or -45.6 ± 3.22, respectively; the mean LN(CHBr_3_ dose in mg/kg DM) for dairy and beef are 2.685 ± 0.1111 and 3.342 ± 0.2099, respectively; and the mean starch content (% DM) for dairy and beef are 12.8 ± 0.22 and 25.7 ± 4.87, respectively. Figure 3 illustrates the observed vs. predicted percentage change in CH_4_ production, yield, and intensity for beef and dairy cattle, using these best-fitting models.

**Figure 3.**
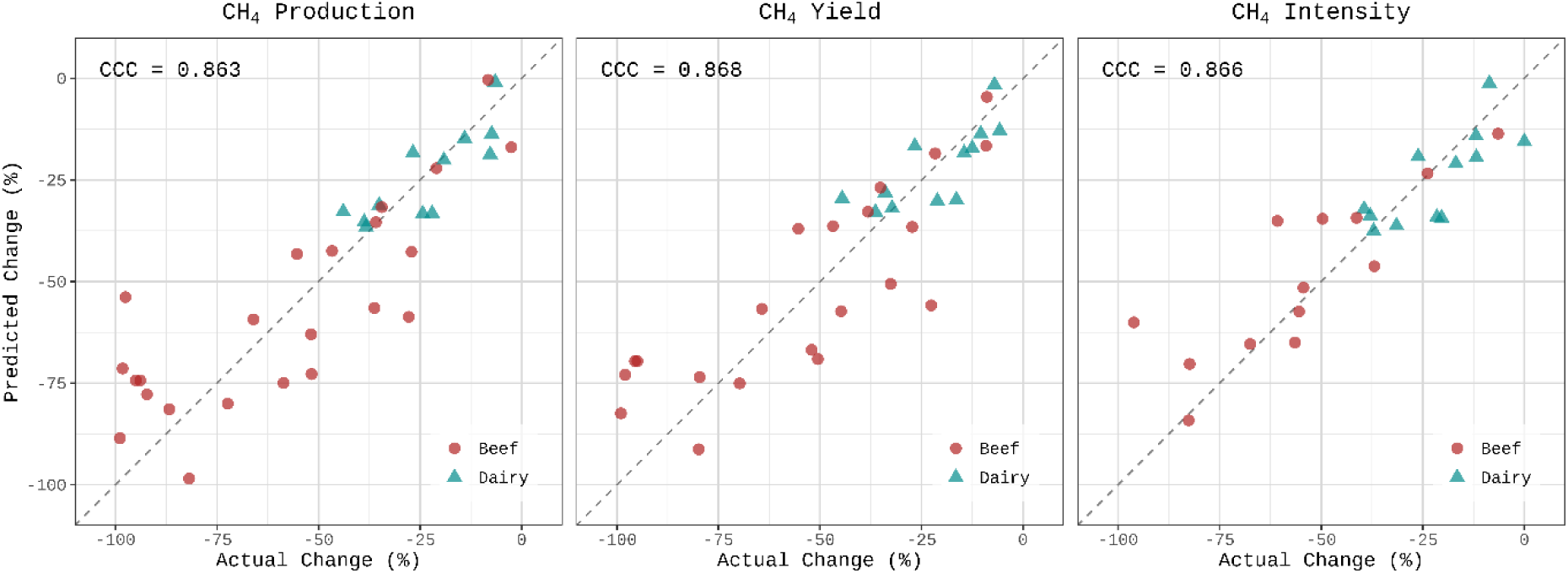
Observed (x-axis) vs. predicted (y-axis) percentage change in CH_4_ production, yield, and intensity for beef (circles) and dairy (triangles) cattle, using the best-fitting models (starch-inclusive models) from Tables 3, 4, and 5 and data without outliers detected by the Cook’s distances method. The dashed diagonal line (y = x) denotes the line of identity, indicating perfect agreement between observed and predicted values. The Concordance Correlation Coefficients (CCC) are presented in the panels of each unit CH_4_ outcome.

All the best-fitting models include starch as a key explanatory variable, reflecting its critical role in shaping CH_4_ responses to CHBr_3_ supplementation. The CCC reported, 0.863 for CH_4_ production, 0.868 for CH_4_ yield, and 0.866 for CH_4_ intensity, demonstrate strong concordance between observed and predicted CH_4_ emissions. In all outcomes, several data points lie near the line of identity, indicating that these starch-inclusive models generally capture both the direction and magnitude of CH_4_ changes across different diets and cattle types. Notably, depending on CH_4_ unit of expression, alternative models that included NDF (for CH_4_ production and yield), FAT (for CH_4_ intensity), and (LN(CHBr_3_ dose))^2^ (when included in models for CH_4_ production and yield) also demonstrated statistically significant effects (*P* ≤ 0.05). These findings indicate that although multiple dietary components contribute to CH_4_ emissions variability, dietary starch is the most robust predictor within our modeling framework, underscoring its importance in refining CH_4_ mitigation strategies in both beef and dairy production systems.

In the beef dataset, the CHBr_3_ dose (mg/kg DM) distribution was left-skewed (Supplemental Figure S1). Due to this asymmetry, a natural log transformation of CHBr_3_ dose before centering was necessary to better represent the data and improve models fit.

Figure 4 visualizes the predicted dose-response relationships from the best-fitting models (incorporating cattle type, natural log-transformed CHBr_3_ dose, and starch content, with the latter two centered at their mean values for each cattle type). The curves demonstrate diminishing returns, where incremental CHBr_3_ additions yield progressively smaller CH_4_ reductions, despite the absence of a quadratic term in those models. This non-linearity arises from the natural log-transformed predictor’s inherent properties: when plotted against the linear x-axis scale (mg/kg DM), the relationship manifests as curvilinear, naturally capturing the biological saturation effect without requiring explicit quadratic terms. Beef cattle showed substantially greater percentage reductions in CH_4_ production and yield than dairy cattle at equivalent doses. However, this difference diminished for CH_4_ intensity, suggesting that emission mitigation effectiveness becomes more comparable between systems when normalized to product output (milk/ECM or ADG). The models predict near-zero or slightly positive relative mitigation differences at very low doses (<3-6 mg/kg DM); however, due to limited data in this range (only 2 observations, at 2 and 6 mg/kg DM), and the theoretical expectation that the inhibitor should not increase CH4 emissions, model predictions at very low doses should be interpreted with caution. The models demonstrate strong CH_4_ reduction potential within practical application ranges (>10 mg/kg DM). Although one might argue that adding a quadratic term on a log-transformed variable could be redundant, our results indicate for CH_4_ production and yield that both the linear and quadratic terms provide unique and important insights. In fact, our frequentist approach revealed that the (LN(CHBr_3_ dose))^2^-NDF-inclusive model (i.e., Type + LN(CHBr_3_ dose) + (LN(CHBr_3_ dose))^2^ + NDF; Tables 3 and 4) significantly represents the variability in CH_4_ production and yield.

**Figure 4.**
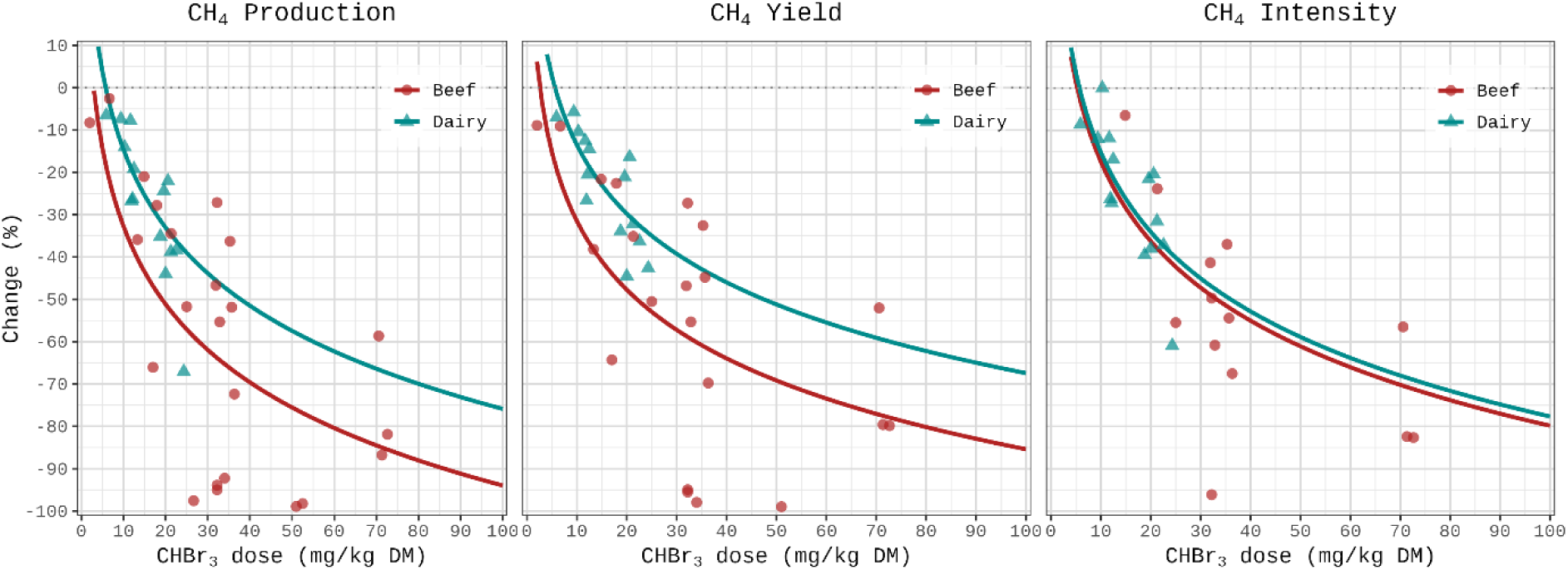
Predicted percentage change (%) in CH_4_ production, CH_4_ yield, and CH_4_ intensity relative to control as a function of CHBr_3_ dose (mg/kg DM). Points represent individual study comparisons for dairy (triangles) and beef (circles). Curves represent predictions for beef (red line) and dairy (cyan line) cattle derived from the best-fitting frequentist meta-regression models (incorporating cattle type, natural log-transformed CHBr_3_ dose, and starch content; see Tables 3-5), holding CHBr_3_ dose and starch at their mean values for each cattle type. Data points do not include outliers detected by the Cook’s distance method.

Across studies, dairy cattle were fed diets with less variation in nutritional composition compared to beef. The nutritional composition of the beef cattle diets varied considerably across studies, with CP ranging from 10.2 to 17.4%, NDF varying between 17.5 and 51.8%, FAT content spanning 3.90 to 8.33%, and starch levels fluctuating from 2.0 to 50.7%. Therefore, the marked variability in beef cattle diets likely underpins the significant impact of dietary covariates in the models, as they capture critical differences in nutritional content that directly affect CH_4_ mitigation outcomes.

The greater CH_4_ mitigation efficacy of CHBr_3_ in high-starch diets compared to high-NDF diets likely arises through interconnected mechanisms centered on hydrogen (H_2_) dynamics and microbial ecology. High-starch fermentation preferentially drives propionate production, a major H_2_ sink, thereby reducing the available H_2_ pool for methanogenesis. In this H_2_-limited environment, CHBr_3_ inhibits a larger proportion of the total methanogenic potential, resulting in greater percentage reductions. In contrast, high-NDF fermentation elevates H_2_ production, creating a larger substrate pool for methanogens and possible larger relative mass of methanogens. Here, while CHBr_3_ maintains similar absolute inhibitory effects, its relative impact diminishes as it acts on a larger potential CH_4_ formation pool. The reduced mitigating effect of CHBr_3_ at higher NDF levels aligns with similar findings for 3-nitrooxypropanol (Dijkstra et al., 2018; Kebreab et al., 2023)

### Publication Bias

Funnel plots (Figure 5) were used to visualize the potential presence of publication bias. These plots display individual study RMDs of CH_4_ emissions on the horizontal axis against their corresponding SEs on the vertical axis. In the random-effects models (i.e., intercept-only model without specifying a mods argument), the central vertical line represents the pooled estimate and the triangular confidence region delineates ±1.96 SE, indicating the range within which 95% of studies would be expected to lie in the absence of bias. A symmetric, inverted funnel shape suggests no substantial publication bias, whereas asymmetry implies that smaller studies might be missing on one side of the funnel, potentially skewing the overall estimate (Afonso et al., 2024). In the random-effects meta-analysis, the funnel plot reveals that many observations fall outside the expected inverted funnel, indicating a high degree of dispersion. This pronounced spread is a hallmark of substantial heterogeneity among the studies, and it may also signal potential small-study effects, such as publication bias or other systematic influences. When moderators (Dairy, Beef, log-centered CHBr_3_ dose, and centered Starch) are subsequently incorporated in a mixed-effects model, the model accounts for some of the between-study variability by adjusting the predicted effect sizes. Consequently, the dispersion observed in the funnel plot is altered; rather than plotting raw effect sizes, the funnel plot often displays standardized residuals that are more closely centered around their predicted values.

**Figure 5.**
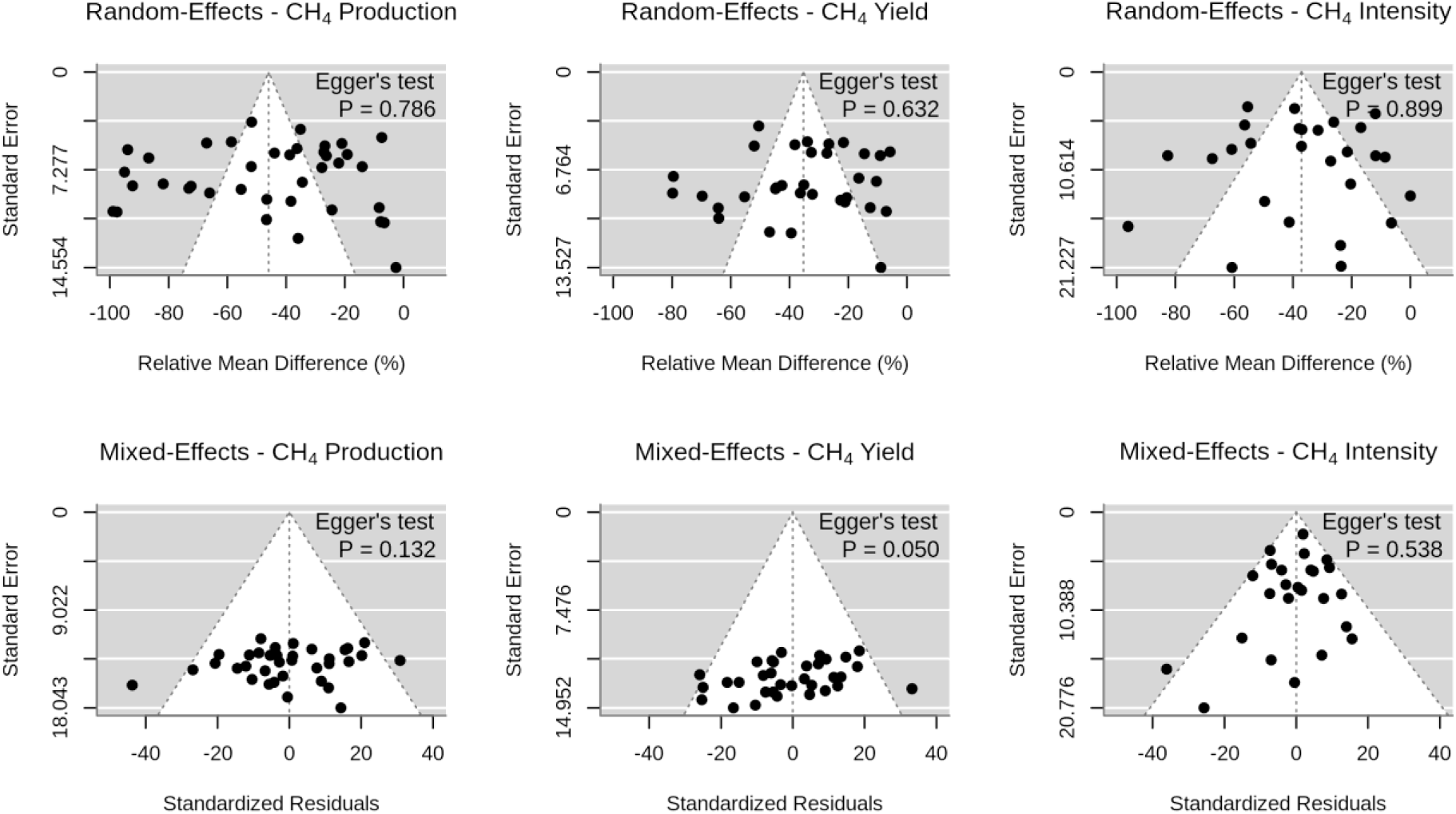
Funnel plots for CH_4_ emissions. Top row: Funnel plots of RMDs vs. SEs for random-effects models of CH_4_ production (g/d), CH_4_ yield (g/kg DMI), and CH_4_ intensity (g/kg milk, ECM, ADG), with pooled estimates and ±1.96 SE regions. Bottom row: Standardized Residuals funnel plots for mixed-effects models, accounting for cattle type (Dairy and Beef), log-centered CHBr_3_ dose and centered starch as moderators. After removing outliers detected with the Cook’s distance method in both models. Symmetry suggests low publication bias. Egger’s test p-values, displayed in each panel, indicate no significant funnel plot asymmetry (*P* ≥ 0.05).

Summaries of the random-effects analyses revealed substantial heterogeneity (*I^2^* > 85%), indicating that the variation in reported RMDs across studies exceeded what would be expected by chance. In the random-effects model *I*^2^ was 94.0, 88.4, and 87.3% for CH_4_ production, yield, and intensity, respectively. In the mixed-effects model (including Dairy, Beef, log-centered CHBr_3_ dose, and centered starch as moderators), *I*^2^ decreased to 76.1, 71.6, and 9.05% for CH_4_ production, yield, and intensity respectively. Notably, the residual heterogeneity for the CH_4_ intensity mixed-effects model dropped substantially (*I*^2^ = 9.05%), indicating that the included moderators (cattle type, log-centered-CHBr3 dose, and starch) explained almost all the between-study variability for this outcome. The resulting standardized residual for the mixed-effects model did not display pronounced asymmetry, suggesting that once moderators were accounted for, the evidence for publication bias diminished. Egger’s regression tests were conducted to statistically evaluate funnel plot asymmetry for each CH_4_ outcome in both random-effects and mixed-effects models. The results indicated no significant (*P* ≥ 0.05) evidence of publication bias across all the CH_4_ outcomes. However, Egger’s test yielded a borderline *P*-value (*P* = 0.050) for the mixed-effects model for CH_4_ yield, suggesting potential slight asymmetry even after accounting for moderators, although visual inspection of the residual plot did not show strong asymmetry. These results suggest that publication bias is unlikely to have substantially influenced the pooled RMD in any of the analyses.

### Bayesian analysis

To ensure a robust and comprehensive analysis, we employed both frequentist and Bayesian statistical approaches. The frequentist meta-regression models allowed us to estimate fixed effects and assess statistical significance using well-established hypothesis testing frameworks. However, frequentist methods have limitations, particularly in handling uncertainty. To address this limitation, we complemented our analysis with a Bayesian framework. Bayesian models allowed us to quantify uncertainty more explicitly through posterior distributions and credible intervals, offering a probabilistic interpretation of the results. By integrating both approaches, we ensured that our conclusions were not solely dependent on *P*-values and frequentist assumptions but were reinforced by Bayesian estimates that account for uncertainty in effect sizes. The comparison of these methodologies provides a more nuanced understanding of how CHBr_3_, DMI, and dietary covariates influence CH_4_ reductions, particularly in cases where frequentist and Bayesian results diverged.

The posterior distributions of the estimated effects on CH_4_ production, yield, and intensity are presented in Figure 6. Each density plot represents the distribution of parameter estimates for different predictors, including (LN(CHBr3 dose))^2^, DMI, starch, FAT, NDF, and CP. The x-axis denotes the estimated effect size and each predictor blue-shaded region illustrates the probability density function of the posterior estimate, with the peak of the curve representing the most probable estimate or the highest density region. The red lines in Figure 6 mark the zero-effect threshold (i.e., no significant effect). If the red line (zero-effect) is outside this highest density region, it indicates stronger evidence for a meaningful estimated effect. If the highest density region is predominantly on one side of this red line, it suggests an effect in that direction (positive or negative). If the peak of the blue curve is very close to the red line, it suggests the effect size is uncertain or weak. The blue shaded areas in Figure 6 also represent the 95% CI, which provide a measure of uncertainty. The symmetric distribution observed in all predictors suggests that the number of Markov chains and iterations were optimal to obtain distributions without strong skewness, which would indicate bias in the estimated effect.

**Figure 6.**
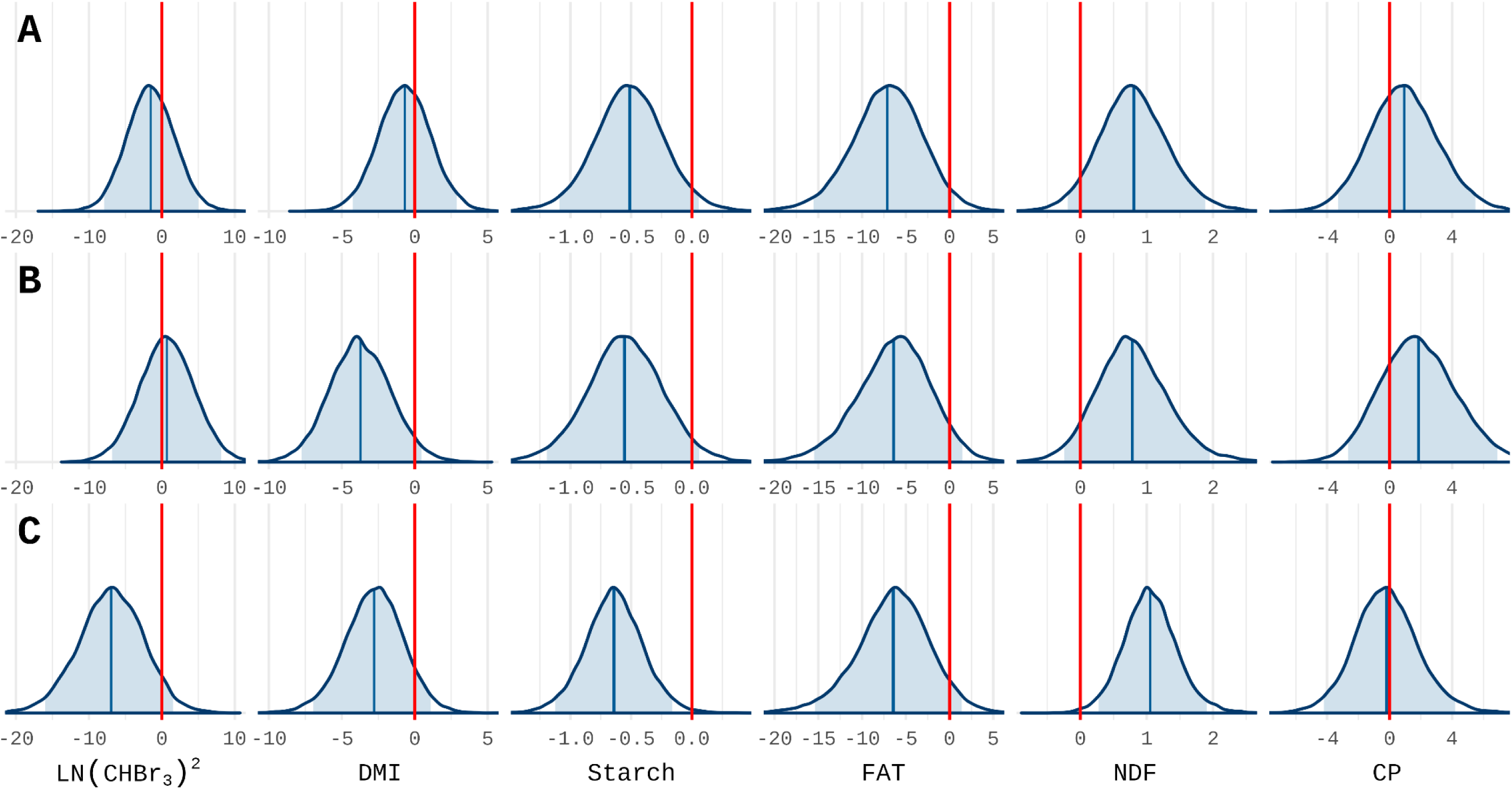
Posterior distributions of covariate effects on CH_4_ production (**A**, g/d), CH_4_ yield (**B**, g/kg DMI), and CH_4_ intensity (**C**, g/kg milk, ECM, ADG). DMI = Dry Matter Intake (kg/day), starch (% DM), FAT = Crude Fat (% DM), NDF = Neutral Detergent Fiber (% DM), and CP = Crude Protein (% DM). Each blue-shaded region illustrates the probability density function of the posterior estimate for a predictor. The peak of each curve represents the most probable estimate (highest density region) of the effect size. Vertical red lines indicate a zero-effect threshold.

Posterior distributions were visually categorized into three effect types: (a) No effect — the zero line lies within the highest density region, indicating no credible effect on CH_4_ reduction; (b) Negative effect — the distribution lies entirely right of zero, suggesting increased CH_4_ emissions despite a positive coefficient; and (c) Positive effect — the distribution lies entirely left of zero, indicating reduced CH_4_ emissions, consistent with a negative and beneficial coefficient.

Visual inspection of the red line’s position relative to the highest density region (peak) of the posterior distribution (Figure 6) reveals that, for CH_4_ production, yield, and intensity, the effects of starch and FAT consistently indicate a meaningful CH_4_-reducing effect (peaks to the left of zero) which allows these factors to be classified in the positive significant effect group. Similarly, DMI shows a meaningful and significant CH_4_-reducing effect for CH_4_ yield, just like DMI and (LN(CHBr3 dose))^2^ for CH_4_ intensity. In contrast, NDF consistently indicates a meaningful CH_4_-increasing effect across all CH_4_ outcomes (peak to the right of zero) which allows this factor to be classified in the negative significant effect group. The other combinations of effects and outcomes can be classified as having no significant effect because the zero-effect line falls within the highest density region of their posterior distributions. This suggests that these covariates do not have a strong or consistent influence on CH_4_ outcomes reduction.

For almost all the covariates, Bayesian and frequentist analyses agreed regarding their significance or lack thereof. However, divergences emerged in the interpretation of (LN(CHBr_3_ dose))^2^, and FAT effects. While the frequentist analysis determined that including (LN(CHBr_3_ dose))^2^ in the model (Dairy or Beef + LN(CHBr_3_ dose) + (LN(CHBr_3_ dose))^2^) was not a significant covariate (*P* > 0.05) for CH_4_ intensity, the most probable values of this effect (i.e., vertical blue line in the highest density region of the posterior distributions; Figure 6) were far from the zero-effect line in CH_4_ intensity, suggesting that there is a significant effect of (LN(CHBr_3_ dose))^2^ on this outcome. While the frequentist analysis determined that FAT was a non-significant covariate for CH_4_ production and yield, the Bayesian analysis indicated greater certainty in the effect of FAT in all CH_4_ outcome reductions, as its highest density region (peak) of the posterior distributions is clearly positioned within the range of negative values. Visually, in Figure 6 (FAT column), the blue shaded 95% CI area is positioned to the left of the zero-effect red line. All Bayesian models achieved strong convergence, as evidenced by Rhat values of 1 and high effective sample sizes, thereby confirming the robustness and reliability of the findings. The observed differences between the Bayesian and frequentist analyses underscore the nuanced ways in which each method handles uncertainty, model structure, and borderline effects, ultimately offering complementary insights rather than mutually exclusive conclusions. In particular, for covariates such as (LN(CHBr_3_ dose))^2^ and FAT, the results differed between the Bayesian and frequentist approaches; suggesting that, for these borderline cases, small shifts in modeling assumptions or data structure can tip the results. However, both methods converged in identifying starch as consistently beneficial (i.e., reducing CH_4_ emission) and NDF as consistently detrimental (i.e., increasing CH_4_ emission), highlighting robust findings for these dietary components.

In applied settings, different analytical methods offer complementary perspectives rather than one being inherently better. Park et al. (2024) used multiple linear regression, k-nearest neighbor, and artificial neural networks to model volatile fatty acid production and CH_4_ emissions, showing that neutral detergent fiber intake was associated with increased acetate and butyrate concentrations, which correlated with higher CH_4_ emissions, while starch intake was linked to lower emissions. Although the k-nearest neighbor method provided the most accurate predictions, the overall agreement across methods highlighted the importance of using multiple approaches to better understand complex biological processes.

### DMI and Product Output

The results of the Random-Effects Meta-Analysis model for DMI and product output (milk, ECM, or ADG) are presented in Figure 7. The frequentist analysis (Table 6) assessed the impact of CHBr_3_ dose on DMI and product output, this analysis revealed that, at mean CHBr_3_ dose (15.8 mg/kg DM), DMI in dairy cattle was significantly (*P* ≤ 0.05) reduced (-6.45%). The reduction in beef cattle (-3.26%) was also significant (*P* ≤ 0.05), at the mean CHBr_3_ dose (37.5 mg/kg DM). Furthermore, the significant negative slope for the CHBr_3_ dose term (-0.269, *P* < 0.001) indicates that increasing the dose relative to these respective means is predicted to further decrease DMI for both cattle types. The model exhibited moderate heterogeneity (*I*^2^ = 39.5%), suggesting variability in DMI was partially explained by cattle type and CHBr_3_ dose. Minor differences were observed depending on the method of outlier removal. When outliers were not removed, the CHBr_3_ dose remained a significant predictor, and DMI reductions in both dairy and beef cattle at the mean CHBr_3_ dose were statistically significant (*P* < 0.05; Supplemental Table S4).

**Figure 7.**
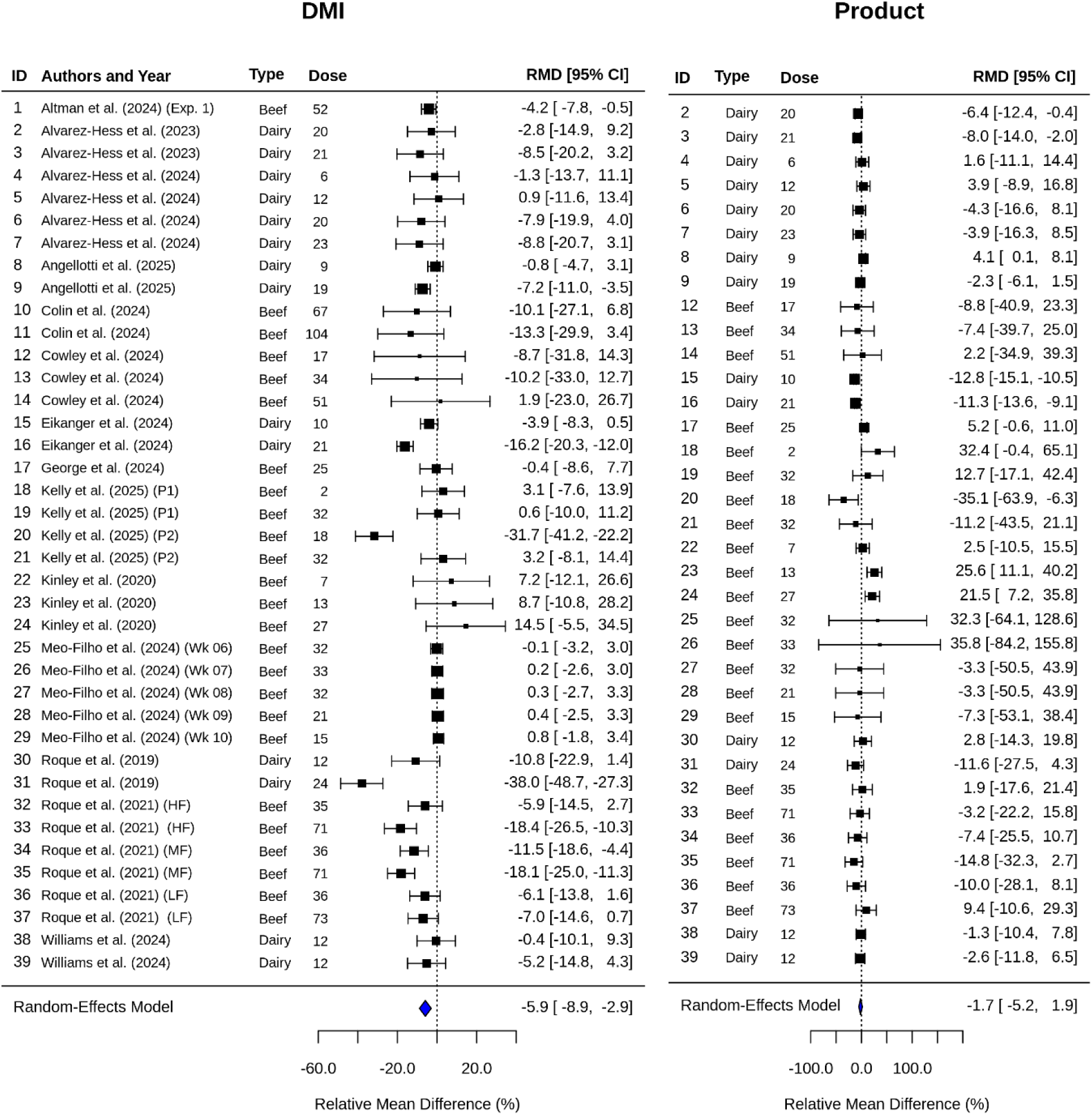
Forest plot showing bromoform (CHBr_3_) dose (mg/kg DM) and relative mean difference values (RMD is calculated as (treatment mean − control treatment mean)/ control treatment mean × 100) in DMI (kg/d) and Products (milk, ECM and ADG, kg/d) for beef and dairy cattle studies. P1 and P2 = experimental periods; Wk = experimental week; HF = high forage diet; LF = low forage diet; MF = medium forage diet. The black squares represent the power of the study (i.e., greater sample sizes and smaller confidence intervals are indicated by a larger box). The summary values at the bottom of the Forest Plot (the diamond, showing - 5.9%, -1.7%) represent the pooled estimate from the Random-Effects Meta-Analysis model.

**Table 6.**
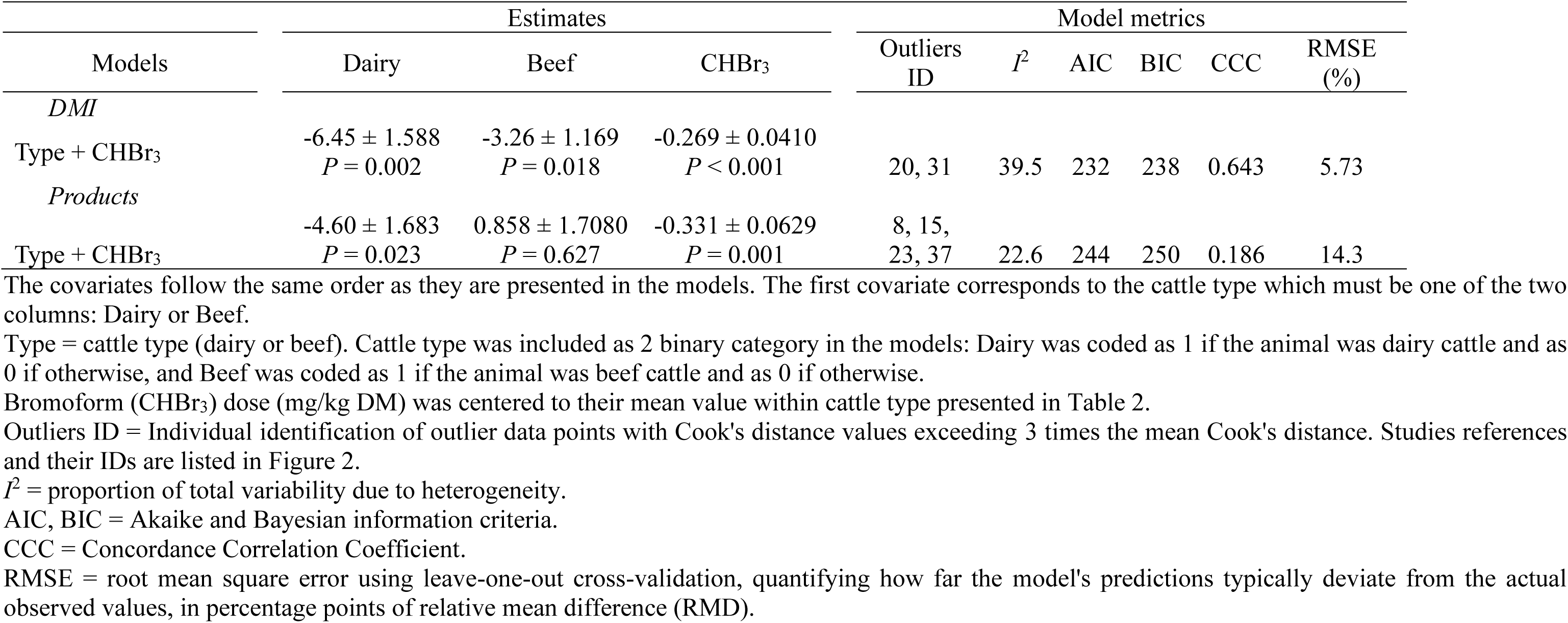
Comparison of regression models evaluating the effect of bromoform dose (CHBr_3_) on the relative mean difference in DMI and products (milk, ECM, ADG) (estimates ± SE)

| Models | Estimates |  |  | Model metrics |  |  |  |  |  |
| --- | --- | --- | --- | --- | --- | --- | --- | --- | --- |
| | Dairy | Beef | CHBr<sub>3</sub> | Outliers ID | $I^2$ | AIC | BIC | CCC | RMSE (%) |
| *DMI* |  |  |  |  |  |  |  |  |  |
| Type + CHBr<sub>3</sub> | -6.45 ± 1.588<br>$P = 0.002$ | -3.26 ± 1.169<br>$P = 0.018$ | -0.269 ± 0.0410<br>$P < 0.001$ | 20, 31 | 39.5 | 232 | 238 | 0.643 | 5.73 |
| *Products* |  |  |  |  |  |  |  |  |  |
| Type + CHBr<sub>3</sub> | -4.60 ± 1.683<br>$P = 0.023$ | 0.858 ± 1.7080<br>$P = 0.627$ | -0.331 ± 0.0629<br>$P = 0.001$ | 8, 15,<br>23, 37 | 22.6 | 244 | 250 | 0.186 | 14.3 |

The covariates follow the same order as they are presented in the models. The first covariate corresponds to the cattle type which must be one of the two columns: Dairy or Beef.
Type = cattle type (dairy or beef). Cattle type was included as 2 binary category in the models: Dairy was coded as 1 if the animal was dairy cattle and as 0 if otherwise, and Beef was coded as 1 if the animal was beef cattle and as 0 if otherwise.
Bromoform (CHBr<sub>3</sub>) dose (mg/kg DM) was centered to their mean value within cattle type presented in Table 2.
$I^2$ = proportion of total variability due to heterogeneity.
AIC, BIC = Akaike and Bayesian information criteria.
CCC = Concordance Correlation Coefficient.
RMSE = root mean square error using leave-one-out cross-validation, quantifying how far the model's predictions typically deviate from the actual observed values, in percentage points of relative mean difference (RMD).

In particular at higher inclusion levels, studies report a significant reduction in DMI, likely due to the cows sorting out the seaweed, reacting to its taste, or related to inflammation of rumen wall. For instance, dairy cows fed *A. taxiformis* at 0.25% of diet DM showed reduced DMI and increased time spent on eating and on eating in a head-down position, suggesting feed sorting or aversion. However, at lower inclusion levels (e.g., 0.125% of diet DM), the reduction in DMI was not present, indicating that palatability challenges may be dose-dependent (Nyløy et al., 2023). Researchers have tried mixing seaweed with molasses or other carriers to mask taste; some success is noted, but at higher doses cattle often still detect the seaweed and reduce intake or sort it out. This underscores that there is an upper limit to how much CHBr_3_ cattle willingly consume. Studies consistently show that cattle do not fully adapt to the taste or presence of *A. taxiformis* in their diet, and in some cases, prolonged supplementation has resulted in significant intake depression. For example, Stefenoni et al. (2021) observed decreased DMI and milk yield in lactating cows fed *A. taxiformis* during 28-day experimental period, while Muizelaar et al. (2021) reported reduced intake across all treatment levels in a 28 day trial. These findings suggest that although *A. taxiformis* is highly effective at reducing enteric CH_4_ emissions, its long-term use may be limited by palatability issues and potential negative effects on feed intake and animal performance.

Notably, dairy cows appear a bit more sensitive to seaweed’s effects on DMI and performance than beef cattle, perhaps because even a moderate intake drop in lactation can noticeably reduce milk outputs. There was a significant negative linear effect (-0.331; *P* ≤ 0.05) of the CHBr_3_ dose on production outcomes (milk, ECM, or ADG). At the average dose of CHBr_3_, a reduction of 4.60% in milk yield was observed (*P* ≤ 0.05), whereas beef cattle showed a non-significant 0.858% increase in ADG. The model exhibited low *I*^2^ (22.6%), indicating that the observed effect of CHBr3 dose on production outcomes was relatively consistent across the different studies included in the analysis, after accounting for cattle type.

Figure 8 illustrates the relationship between CHBr_3_ dose and the RMD in DMI and products (milk, ECM, or ADG). The superimposed lines represent predicted linear dose-responses from meta-regression models after exluding outliers identified by Cook’s distance. Several data points indicated positive RMDs (i.e., increased ADG) including low doses (e.g., ID 18: +32.4% ADG at 2 mg/kg) and moderate-to-higher dose responses (e.g., ID 24: +21.5% at 27 mg/kg; ID 25: +32.3% at 32 mg/kg; ID 26: +35.8% at 33 mg/kg). However, these points were not flagged as outliers by Cook’s distance method. The presence of these variable responses contributed to poor model fit (CCC = 0.186) suggesting that a linear model may inadequately capture the ADG-dose relationship in beef cattle. Biological complexity or data noise could underlie this incosistency. Indeed, a negative RMD (i.e., decreased ADG) occurred, with the lowest response being -35.1% (ID 20, 18 mg/kg DM), but this data point was not flagged as an outlier by Cook’s distance method either. The inconsistent response (both increased and decreased ADG within a range of 2 to 35 mg/kg of CHBr_3_) does not enable a model to adequately capture variation in response. When no outliers were removed, at the average dose of CHBr_3_, dairy cattle showed a non-significant decrease in milk production, and beef cattle showed a non-significant increase in ADG (Supplemental Table S4). An alternative outlier removal approach (CCC method, Supplemental Table S8) excluded IDs 18, 20, 25, and 26 (i.e., the changes in ADG with the greatest decrease (ID 20) and greatest increases (ID 18, 25, and 26)), modestly improving predictive metrics (CCC: 0.280; RMSE: 10.2%) compared to the Cook’s distance approach.

**Figure 8.**
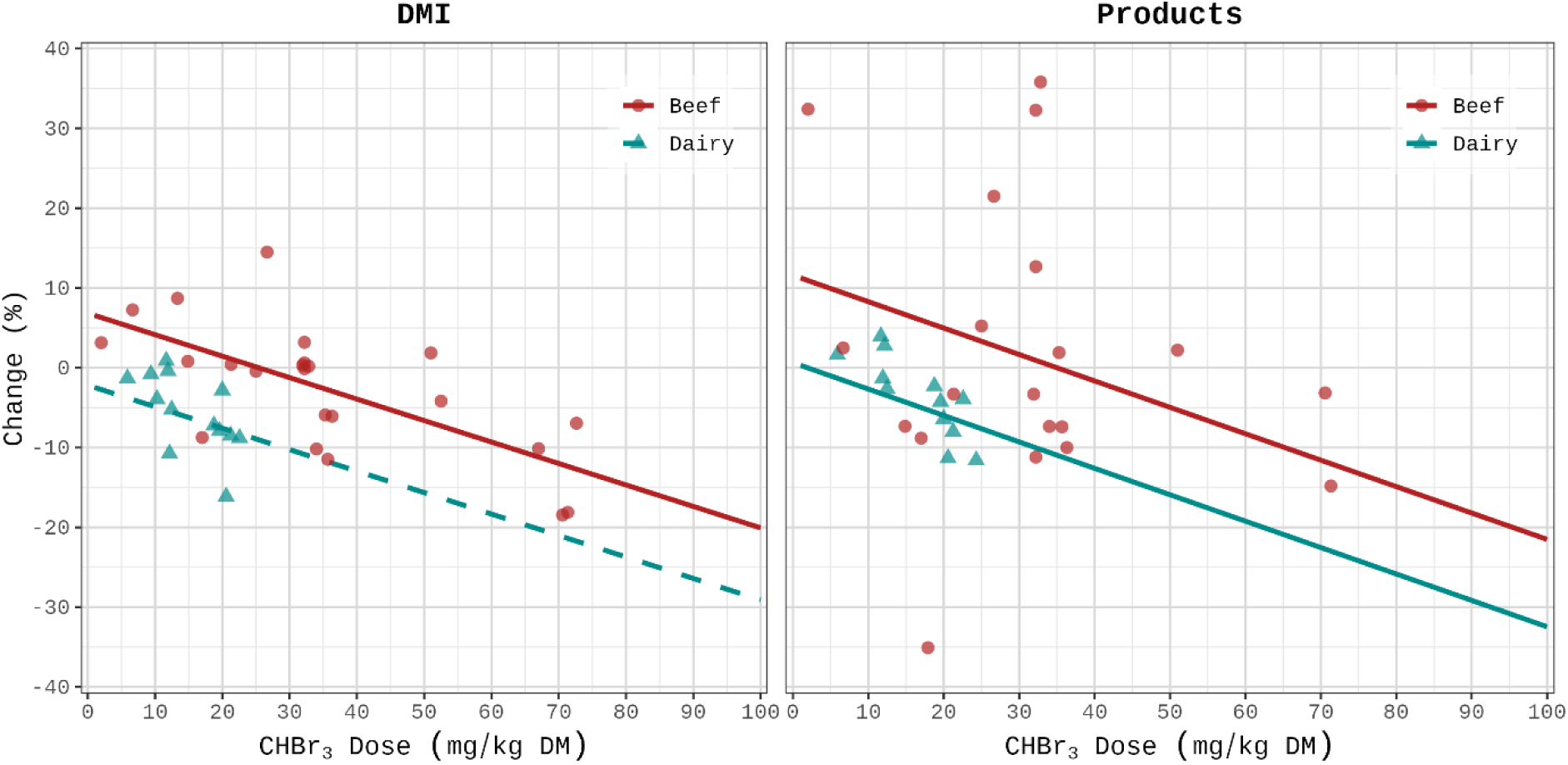
Predicted percentage change (%) in DMI and products (milk, ECM, or ADG) relative to control as a function of CHBr_3_ dose (mg/kg DM). Points represent individual study comparisons for dairy (triangles) and beef (circles). Lines represent predictions for beef (red line) and dairy (cyan line) cattle derived from the best-fitting frequentist meta-regression models (incorporating cattle type and non-log-transformed CHBr_3_ dose; see Table 6). Data points do not include outliers detected by the Cook’s distance method.

### Bromoform Carrier, Methane Measurement Technique, and Cattle Type

Table 7 summarizes the estimated RMD differences in CH_4_ production, yield, and intensity across carriers, measurement techniques, and cattle types.

**Table 7.** Estimated difference in relative mean difference for CH_4_ production (g/d), CH_4_ yield (g/kg DMI), and CH_4_ intensity (g/kg milk, ECM, ADG) across carriers, techniques, and cattle types using a mixed-effects meta-regression model incorporating log-centered-CHBr_3_ dose (mg/kg DM)

| Contrasts | CH<sub>4</sub> Production |  |  | CH<sub>4</sub> Yield |  |  | CH<sub>4</sub> Intensity |  |  |
| --- | --- | --- | --- | --- | --- | --- | --- | --- | --- |
|  | Estimate<sup>1</sup> | SE | P | Estimate | SE | P | Estimate | SE | P |
| *Carrier* |  |  |  |  |  |  |  |  |  |
| Biomass - Oil | 21.5 | 11.9 | 0.081 | 22.4 | 11.4 | 0.060 | 24.7 | 11.1 | 0.036 |
| *Technique* |  |  |  |  |  |  |  |  |  |
| GF - RC | 22.3 | 12.8 | 0.207 | 25.9 | 12.2 | 0.104 | NA | NA | NA |
| GF - SF<sub>6</sub> | -24.7 | 13.7 | 0.188 | -19.8 | 10.5 | 0.164 | -31.6 | 11.5 | 0.011 |
| RC - SF<sub>6</sub> | -47.0 | 13.5 | 0.005 | -45.7 | 10.3 | < 0.001 | NA | NA | NA |
| *Cattle Type* |  |  |  |  |  |  |  |  |  |
| Dairy - Beef | 14.2 | 9.33 | 0.140 | 20.1 | 4.41 | < 0.001 | 17.7 | 5.44 | 0.004 |

<sup>1</sup>Estimate refers to the difference in relative mean difference (% RMD). A positive estimate signifies that the first group in the contrast has a higher RMD (less reduction) compared to the second group. Conversely, a negative estimate indicates the first group has a lower RMD (greater reduction).
Biomass = Predominantly seaweed-based carriers or algae-derived manufactured products. Oil = Edible oil carriers (with synthetic CHBr<sub>3</sub> or seaweed biomass). GF = GreenFeed system, RC = Respiration Chamber, SF<sub>6</sub> = Sulfur hexafluoride tracer gas technique.

Biomass-based studies used higher CHBr_3_ doses (29.8–32.4 mg/kg DM) than oil-based studies (19.5–22.6 mg/kg DM). Despite this, oil was associated with greater CH_4_ reductions, significantly so for CH_4_ intensity (*P* < 0.05), and nearing significance for production and yield (*P* < 0.10), suggesting carrier type influences efficacy.

Average CHBr_3_ doses varied by measurement technique: GreenFeed studies used the highest dose (33.8 mg/kg DM), followed by respiration chamber (24.8 mg/kg DM for CH_4_ production and yield), and SF_6_ (15.6 mg/kg DM). Respiration chamber was associated with significantly greater CH_4_ reductions than SF_6_ for production and yield (*P* < 0.05). Differences between GreenFeed and the other techniques were not significant for CH_4_ production or yield (*P* ≥ 0.05). For CH_4_ intensity, GreenFeed resulted in significantly greater reductions compared to SF_6_ (*P* < 0.05); data were insufficient for comparisons involving respiration chamber. Although GreenFeed studies used higher doses, adjusted CH_4_ reductions were not greater than those observed with respiration chamber, suggesting that measurement technique may affect outcomes beyond dose differences alone.

Beef cattle received a substantially higher average CHBr_3_ dose than dairy cattle (34.6 vs. 15.8 mg/kg DM for CH_4_ production and yield; 38.9 vs. 15.8 mg/kg DM for CH_4_ intensity). After adjusting for dose, carrier, and measurement technique, RMD was significantly lower (*P* < 0.05) in beef cattle for CH_4_ yield and intensity, indicating greater reductions. No significant difference (*P* ≥ 0.05) was observed for CH_4_ production between cattle types.

## CONCLUSIONS

The study confirms that CHBr_3_-containing seaweeds and CHBr_3_-based additives significantly reduce CH_4_ emission in dairy and beef cattle. The pooled estimates from the Random-Effects Meta-Analysis model show reductions of 47.3, 43.3, and 39.0% for CH_4_ production, yield, and intensity, respectively, at average CHBr_3_ dose level of ≈28.3 mg/kg DM, with increases in CHBr_3_ dose resulting in larger efficacy in mitigating CH_4_ emissions. The Bayesian framework provided critical confirmation of dietary starch and NDF effects. For starch, the entire posterior density for all CH_4_ outcomes fell entirely to the left of the zero-effect threshold, indicating a high probability of its beneficial role in enhancing CHBr_3_ efficacy. Conversely, NDF’s posterior density resided entirely to the right of zero, reflecting its consistent association with diminished CHBr_3_ effectiveness. This Bayesian confirmation aligns with frequentist meta-regression results, solidifying starch and NDF as robust, directionally consistent predictors of CH_4_ mitigation outcomes. At their respective average CHBr_3_ doses, DMI was significantly reduced in dairy and beef cattle. Similarly, product output at the average dose was significantly reduced for dairy. The results highlight the potential for optimizing dietary management to maximize the environmental benefits of CHBr_3_. The detailed predictive equations provide a basis to integrate these findings into GHG accounting systems and practical farm management strategies.

## NOTES

The study was financially supported by the UC Davis Sesnon Endowed Chair program (EK and JFRA). Special thanks to Robert Kinley, Breanna Roque, Leanna Kelly and Paulo Meo-Filho for their generous provision of additional data, which greatly enhanced the scope and depth of our research.

## Nonstandard abbreviations used

CHBr_3_: bromoform
CH_4_: methane
CO_2_: carbon dioxide
RC: respiration chamber
GF: GreenFeed
*I*^2^: heterogeneity
LOOCV: leave-one-out cross-validation
MD: mean difference
RMD: relative mean difference
RMSE: root mean square error
SF_6_: sulfur hexafluoride
STP: standard temperature and pressure

## Notes

### Competing Interest Statement

The authors have declared no competing interest.

