## Supplemental Information for "A meta-analysis of effects of seaweed and other bromoform containing feed ingredients on methane production, yield, and intensity in cattle"

**Figure S1.**

Supplemental Figure S1 presents the distribution of bromoform (CHBr_3_) dose before the application of the log transformation. The figure highlights the significant skewness in the raw CHBr_3_ dose values, particularly for beef cattle, as confirmed by a Shapiro-Wilk test *P*-value of 0.031. The log transformation was a critical preprocessing step, as it addressed the skewness and improved the suitability of the data for subsequent analyses. By normalizing the distribution, the transformation facilitated the interpretation of model results. Importantly, all equations derived from the analysis assume that the input CHBr_3_ doses are log-transformed. Using untransformed data in these equations would yield incorrect results, as the relationships were defined on the log scale.

**Table S1 to Table S4.** **No outlier elimination.**

Tables S2 to S4 present the results of models evaluating the effects of CHBr_3_ dose and additional dietary or animal covariates on methane emission metrics (CH_4_ production, yield, and intensity) and on DMI and products outputs (milk, ECM, or ADG) without outlier removal. The tables include *P*-values to assess the statistical significance of each covariate's effect.

**Table S5 to Table S8.**  **Outlier elimination based on concordance correlation coefficient (CCC).**

Tables S5 to S8 present the results of models after outlier elimination using the Concordance Correlation Coefficient (CCC)-based method. Our CCC-based method was adopted, focusing on prediction accuracy rather than coefficient stability. This approach iteratively removes observations with the largest residuals (discrepancies between predicted and actual values of the response variable), refits the model, and recalculates predictions until the number of removed points matches those identified by the Cook’s distance method. By prioritizing prediction accuracy and using the CCC to evaluate model performance, the CCC-based method provided a balanced, data-driven criterion for outlier removal.

**Example: Calculating change (%) in CH_4_ production**

This example demonstrates how to calculate the percentage change in CH_4_ production for a dairy cow based on specific characteristics, including the dose of CHBr_3_, and dietary starch content.

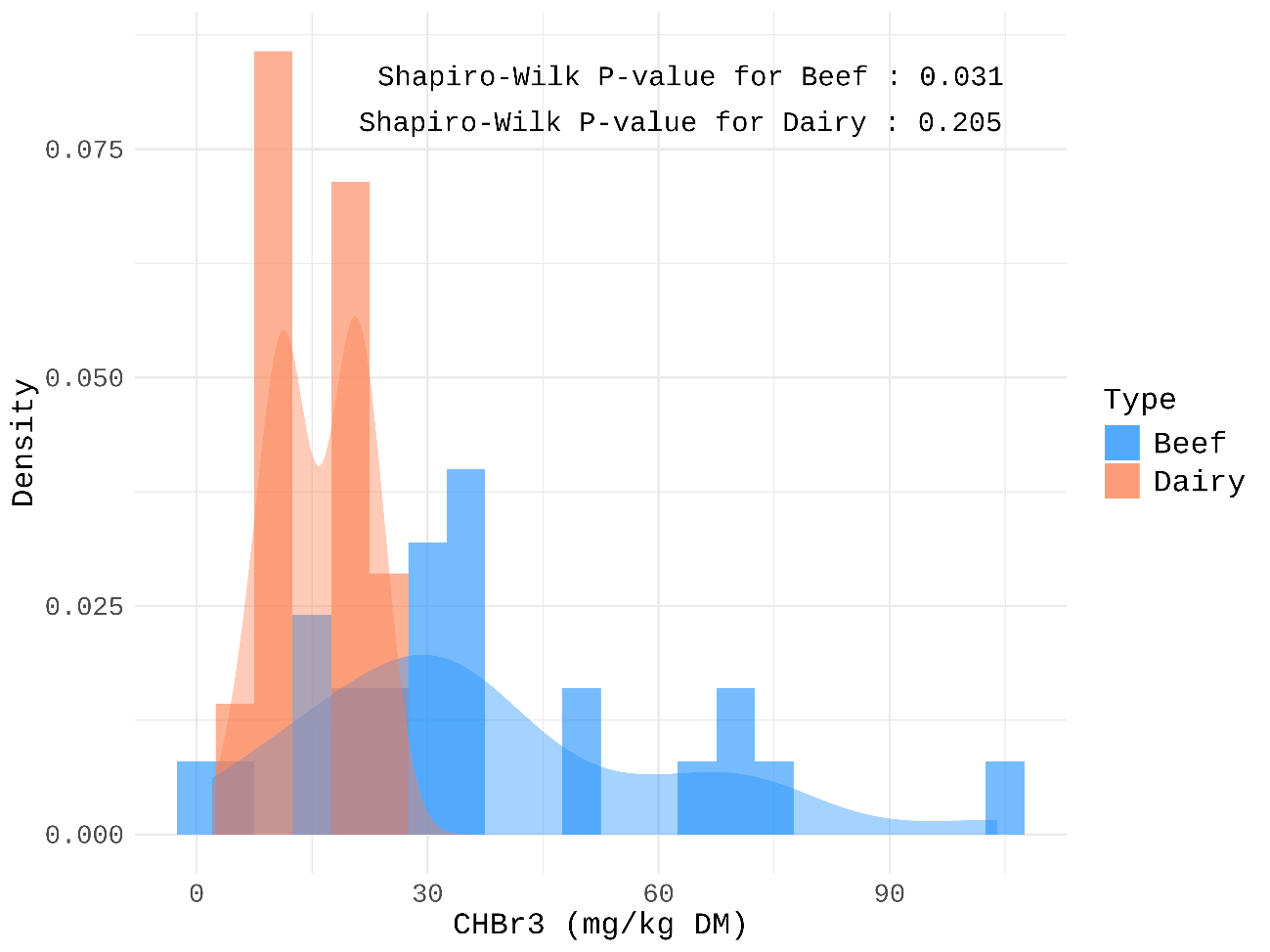

### Figure S1. Histogram of CHBr_3_ dose (mg/kg DM) by cattle type

### **Table S1.** Comparison of regression models evaluating the effect of bromoform (CHBr_3_; in LN mg CHBr_3_/kg DM) dose and additional dietary or animal covariates on the relative mean difference in CH_4_ production (estimates ± SE) without outliers removal

|  |  | Estimates | | | |  | Model metrics | | | | |
| --- | --- | --- | --- | --- | --- | --- | --- | --- | --- | --- | --- |
| Models |  | Dairy | Beef | 2nd  Covariate | 3rd  Covariate |  | *I*^2^ | AIC | BIC | CCC | RMSE (%) |
| Type + CHBr_3_ |  | -27.4 ± 3.63 *P* < 0.001 | -58.1 ± 7.00 *P* < 0.001 | -23.9 ± 3.90 *P* < 0.001 | NA |  | 85.9 | 348 | 355 | 0.737 | 20.3 |
| Type + CHBr_3_ + (CHBr_3_)^2^ |  | -27.3 ± 3.76 *P* < 0.001 | -57.8 ± 8.29 *P* < 0.001 | -24.4 ± 6.58 *P* = 0.004 | -0.489 ± 3.6733 *P* = 0.897 |  | 86.0 | 350 | 358 | 0.737 | 26.1 |
| Type + CHBr_3_ + DMI |  | -27.4 ± 3.89 *P* < 0.001 | -58.2 ± 7.26 *P* < 0.001 | -23.7 ± 3.84 *P* < 0.001 | 0.624 ± 2.0227 *P* = 0.764 |  | 85.7 | 350 | 358 | 0.738 | 21.0 |
| Type + CHBr_3_ + CP |  | -27.4 ± 3.96 *P* < 0.001 | -58.1 ± 7.00 *P* < 0.001 | -24.0 ± 4.11 *P* < 0.001 | -0.975 ± 2.9171 *P* = 0.745 |  | 85.7 | 349 | 358 | 0.740 | 20.9 |
| Type + CHBr_3_ + NDF |  | -27.5 ± 3.41 *P* < 0.001 | -58.1 ± 5.77 *P* < 0.001 | -22.7 ± 4.47 *P* < 0.001 | 0.714 ± 0.3029 *P* = 0.040 |  | 83.3 | 345 | 353 | 0.773 | 19.4 |
| Type + CHBr_3_ + Starch |  | -24.6 ± 2.65 *P* < 0.001 | -58.1 ± 4.89 *P* < 0.001 | -22.0 ± 4.52 *P* = 0.001 | -0.552 ± 0.2573 *P* = 0.061 |  | 81.1 | 324 | 332 | 0.801 | 18.8 |
| Type + CHBr_3_ + FAT |  | -27.2 ± 3.68 *P* < 0.001 | -58.4 ± 6.98 *P* < 0.001 | -24.4 ± 5.02 *P* = 0.001 | -4.331 ± 2.8282 *P* = 0.157 |  | 84.6 | 347 | 355 | 0.757 | 20.8 |

The covariates follow the same order as they are presented in the models. The first covariate corresponds to the cattle type which must be one of the two columns: Dairy or Beef. For example, in the model 'Type + CHBr_3_ + Starch', the '2nd Covariate' column shows the estimate for the (log-transformed, centered) CHBr_3_ dose, and the '3rd Covariate' column shows the estimate for (centered) Starch.

Type = cattle type (dairy or beef). Cattle type was included as 2 binary category in the models: Dairy was coded as 1 if the animal was dairy cattle and as 0 if otherwise, and Beef was coded as 1 if the animal was beef cattle and as 0 if otherwise.

All variables (except cattle type) were centered to their mean value within each cattle type presented in Table 1 in the main document.

Bromoform (CHBr_3_) dose (mg/kg DM) was natural log-transformed before models fitting.

*I*^2^ = proportion of total variability due to heterogeneity.

AIC, BIC = Akaike and Bayesian information criteria.

CCC = Concordance Correlation Coefficient.

RMSE = root mean square error using leave-one-out cross-validation, quantifying how far the model's predictions typically deviate from the actual observed values, in percentage points of relative mean difference (RMD).

### **Table S2.** Comparison of regression models evaluating the effect of bromoform (CHBr_3_; in LN mg CHBr3/kg DM) dose and additional dietary or animal covariates on the relative mean difference in CH_4_ yield (estimates ± SE) without outliers removal

|  |  | Estimates | | | |  | Model metrics | | | | |
| --- | --- | --- | --- | --- | --- | --- | --- | --- | --- | --- | --- |
| Models |  | Dairy | Beef | 2nd  Covariate | 3rd  Covariate |  | *I*^2^ | AIC | BIC | CCC | RMSE (%) |
| Type + CHBr_3_ |  | -23.2 ± 2.30 *P* < 0.001 | -54.9 ± 7.07 *P* < 0.001 | -20.5 ± 3.85 *P* < 0.001 | NA |  | 86.8 | 341 | 347 | 0.703 | 20.5 |
| Type + CHBr_3_ + (CHBr_3_)^2^ |  | -23.5 ± 2.52 *P* < 0.001 | -56.0 ± 8.50 *P* < 0.001 | -19.0 ± 5.49 *P* = 0.007 | 1.71 ± 3.828 *P* = 0.666 |  | 86.8 | 342 | 351 | 0.705 | 25.1 |
| Type + CHBr_3_ + DMI |  | -23.1 ± 2.21 *P* < 0.001 | -55.0 ± 7.64 *P* < 0.001 | -20.6 ± 3.96 *P* = 0.001 | -1.75 ± 1.959 *P* = 0.395 |  | 86.4 | 342 | 350 | 0.712 | 20.6 |
| Type + CHBr_3_ + CP |  | -23.2 ± 2.44 *P* < 0.001 | -54.9 ± 7.40 *P* < 0.001 | -20.5 ± 4.26 *P* = 0.001 | -0.290 ± 3.0765 *P* = 0.927 |  | 86.7 | 343 | 351 | 0.703 | 21.3 |
| Type + CHBr_3_ + NDF |  | -23.2 ± 2.39 *P* < 0.001 | -55.1 ± 6.22 *P* < 0.001 | -19.1 ± 4.93 *P* = 0.004 | 0.654 ± 0.2725 *P* = 0.040 |  | 84.9 | 338 | 347 | 0.737 | 20.0 |
| Type + CHBr_3_ + Starch |  | -22.3 ± 2.66 *P* < 0.001 | -55.4 ± 5.52 *P* < 0.001 | -18.8 ± 5.14 *P* = 0.006 | -0.560 ± 0.2674 *P* = 0.070 |  | 84.6 | 320 | 328 | 0.759 | 19.9 |
| Type + CHBr_3_ + FAT |  | -23.1 ± 2.97 *P* < 0.001 | -54.7 ± 7.29 *P* < 0.001 | -21.5 ± 5.05 *P* = 0.002 | -3.56 ± 2.414 *P* = 0.174 |  | 85.7 | 341 | 349 | 0.720 | 21.0 |

The covariates follow the same order as they are presented in the models. The first covariate corresponds to the cattle type which must be one of the two columns: Dairy or Beef. For example, in the model 'Type + CHBr_3_ + Starch', the '2nd Covariate' column shows the estimate for the (log-transformed, centered) CHBr_3_ dose, and the '3rd Covariate' column shows the estimate for (centered) Starch.

Type = cattle type (dairy or beef). Cattle type was included as 2 binary category in the models: Dairy was coded as 1 if the animal was dairy cattle and as 0 if otherwise, and Beef was coded as 1 if the animal was beef cattle and as 0 if otherwise.

All variables (except cattle type) were centered to their mean value within each cattle type presented in Table 1 in the main document.

Bromoform (CHBr_3_) dose (mg/kg DM) was natural log-transformed before models fitting.

*I*^2^ = proportion of total variability due to heterogeneity.

AIC, BIC = Akaike and Bayesian information criteria.

CCC = Concordance Correlation Coefficient.

RMSE = root mean square error using leave-one-out cross-validation, quantifying how far the model's predictions typically deviate from the actual observed values, in percentage points of relative mean difference (RMD).

### **Table S3.** Comparison of regression models evaluating the effect of bromoform (CHBr_3_; in LN mg CHBr_3_/kg DM) dose and additional dietary or animal covariates on the relative mean difference in CH_4_ intensity (estimates ± SE) without outliers removal

|  |  | Estimates | | | |  | Model metrics | | | | |
| --- | --- | --- | --- | --- | --- | --- | --- | --- | --- | --- | --- |
| Models |  | Dairy | Beef | 2nd  Covariate | 3rd  Covariate |  | *I*^2^ | AIC | BIC | CCC | RMSE (%) |
| Type + CHBr_3_ |  | -25.9 ± 3.11 *P* < 0.001 | -50.7 ± 5.67 *P* < 0.001 | -24.6 ± 4.61 *P* = 0.001 | NA |  | 69.4 | 266 | 272 | 0.648 | 23.8 |
| Type + CHBr_3_ + (CHBr_3_)^2^ |  | -24.9 ± 3.30 *P* < 0.001 | -46.7 ± 5.41 *P* < 0.001 | -29.5 ± 6.06 *P* = 0.003 | -6.47 ± 2.604 *P* = 0.048 |  | 69.4 | 266 | 273 | 0.689 | 29.3 |
| Type + CHBr_3_ + DMI |  | -25.5 ± 3.94 *P* = 0.001 | -49.8 ± 6.26 *P* < 0.001 | -26.3 ± 4.79 *P* = 0.002 | -2.21 ± 3.159 *P* = 0.511 |  | 68.5 | 267 | 274 | 0.680 | 25.2 |
| Type + CHBr_3_ + CP |  | -25.9 ± 3.46 *P* < 0.001 | -50.3 ± 5.34 *P* < 0.001 | -24.4 ± 4.56 *P* = 0.002 | -0.704 ± 2.3174 *P* = 0.772 |  | 70.1 | 268 | 275 | 0.654 | 24.2 |
| Type + CHBr_3_ + NDF |  | -26.5 ± 2.80 *P* < 0.001 | -47.5 ± 5.15 *P* < 0.001 | -23.8 ± 2.68 *P* < 0.001 | 0.937 ± 0.3683 *P* = 0.044 |  | 49.5 | 260 | 267 | 0.704 | 21.6 |
| Type + CHBr_3_ + Starch |  | -24.6 ± 2.22 *P* < 0.001 | -48.1 ± 3.20 *P* < 0.001 | -24.7 ± 2.21 *P* < 0.001 | -0.632 ± 0.1773 *P* = 0.016 |  | 41.2 | 242 | 248 | 0.716 | 21.5 |
| Type + CHBr_3_ + FAT |  | -26.1 ± 2.95 *P* < 0.001 | -49.2 ± 6.74 *P* < 0.001 | -23.2 ± 4.45 *P* = 0.002 | -4.897 ± 2.8045 *P* = 0.131 |  | 62.6 | 266 | 273 | 0.669 | 23.2 |

The covariates follow the same order as they are presented in the models. The first covariate corresponds to the cattle type which must be one of the two columns: Dairy or Beef. For example, in the model 'Type + CHBr_3_ + Starch', the '2nd Covariate' column shows the estimate for the (log-transformed, centered) CHBr_3_ dose, and the '3rd Covariate' column shows the estimate for (centered) Starch.

Type = cattle type (dairy or beef). Cattle type was included as 2 binary category in the models: Dairy was coded as 1 if the animal was dairy cattle and as 0 if otherwise, and Beef was coded as 1 if the animal was beef cattle and as 0 if otherwise.

All variables (except cattle type) were centered to their mean value within each cattle type presented in Table 1 in the main document.

Bromoform (CHBr_3_) dose (mg/kg DM) was natural log-transformed before models fitting.

*I*^2^ = proportion of total variability due to heterogeneity.

AIC, BIC = Akaike and Bayesian information criteria.

CCC = Concordance Correlation Coefficient.

RMSE = root mean square error using leave-one-out cross-validation, quantifying how far the model's predictions typically deviate from the actual observed values, in percentage points of relative mean difference (RMD).

### **Table S4.** Comparison of regression models evaluating the effect of bromoform (CHBr_3_) dose on the relative mean difference in DMI and products (milk, ECM, ADG) (estimates ± SE) without outliers removal

|  |  | Estimates | | |  | Model metrics | | | | |
| --- | --- | --- | --- | --- | --- | --- | --- | --- | --- | --- |
| Models |  | Dairy | Beef | CHBr_3_ |  | *I*^2^ | AIC | BIC | CCC | RMSE (%) |
| *DMI* |  |  |  |  |  |  |  |  |  |  |
| Type + CHBr_3_ |  | -7.92 ± 2.721 *P* = 0.014 | -4.69 ± 1.804 *P* = 0.025 | -0.223 ± 0.0735 *P* = 0.011 |  | 81.3 | 286 | 293 | 0.347 | 9.35 |
| *Products* |  |  |  |  |  |  |  |  |  |  |
| Type + CHBr_3_ |  | -4.61 ± 2.533 *P* = 0.102 | 3.09 ± 2.842 *P* = 0.305 | -0.263 ± 0.0727 *P* = 0.006 |  | 58.1 | 282 | 289 | 0.200 | 14.5 |

The covariates follow the same order as they are presented in the models. The first covariate corresponds to the cattle type which must be one of the two columns: Dairy or Beef.

Type = cattle type (dairy or beef). Cattle type was included as 2 binary category in the models: Dairy was coded as 1 if the animal was dairy cattle and as 0 if otherwise, and Beef was coded as 1 if the animal was beef cattle and as 0 if otherwise.

Bromoform (CHBr_3_) dose (mg/kg DM) was centered to their mean value within cattle type presented in Table 2 in the main document.

Outliers ID = Individual identification of outlier data points using the CCC-based method. Studies references and their IDs are listed in Figure 2 in the main document.

*I*^2^ = proportion of total variability due to heterogeneity.

AIC, BIC = Akaike and Bayesian information criteria.

CCC = Concordance Correlation Coefficient.

RMSE = root mean square error using leave-one-out cross-validation, quantifying how far the model's predictions typically deviate from the actual observed values, in percentage points of relative mean difference (RMD).

### **Table S5.** Comparison of regression models evaluating the effect of bromoform (CHBr_3_; in LN mg CHBr_3_/kg DM) dose and additional dietary or animal covariates on the relative mean difference in CH_4_ production (estimates ± SE) with identification of outliers using the CCC-based method

|  |  | Estimates | | | |  | Model metrics | | | | | |
| --- | --- | --- | --- | --- | --- | --- | --- | --- | --- | --- | --- | --- |
| Models |  | Dairy | Beef | 2nd  Covariate | 3rd  Covariate |  | Outliers  ID | *I*^2^ | AIC | BIC | CCC | RMSE (%) |
| Type + CHBr_3_ |  | -27.4 ± 3.63 *P* < 0.001 | -54.9 ± 6.57 *P* < 0.001 | -24.1 ± 3.83 *P* < 0.001 | NA |  | 19, 24 | 83.3 | 323 | 330 | 0.766 | 18.7 |
| Type + CHBr_3_ + (CHBr_3_)^2^ |  | -27.1 ± 3.70 *P* < 0.001 | -53.4 ± 7.82 *P* < 0.001 | -26.1 ± 6.25 *P* = 0.002 | -2.16 ± 3.407 *P* = 0.541 |  | 19, 24 | 83.2 | 325 | 333 | 0.769 | 22.7 |
| Type + CHBr_3_ + DMI |  | -27.6 ± 3.95 *P* < 0.001 | -52.9 ± 6.57 *P* < 0.001 | -23.6 ± 3.82 *P* < 0.001 | 0.922 ± 1.8763 *P* = 0.634 |  | 19, 21, 24 | 79.8 | 312 | 320 | 0.779 | 18.8 |
| Type + CHBr_3_ + CP |  | -27.4 ± 3.83 *P* < 0.001 | -54.9 ± 6.66 *P* < 0.001 | -24.1 ± 3.98 *P* < 0.001 | -0.196 ± 2.7004 *P* = 0.944 |  | 19, 24 | 83.3 | 325 | 333 | 0.766 | 19.5 |
| Type + CHBr_3_ + NDF |  | -27.7 ± 3.32 *P* < 0.001 | -56.7 ± 5.83 *P* < 0.001 | -23.3 ± 3.57 *P* < 0.001 | 0.901 ± 0.1970 *P* = 0.001 |  | 1, 10, 24 | 77.8 | 307 | 315 | 0.832 | 16.6 |
| Type + CHBr_3_ + Starch |  | -24.8 ± 2.55 *P* < 0.001 | -58.0 ± 4.31 *P* < 0.001 | -24.1 ± 3.48 *P* < 0.001 | -0.692 ± 0.1845 *P* = 0.005 |  | 10, 24 | 74.6 | 295 | 303 | 0.870 | 15.8 |
| Type + CHBr_3_ + FAT |  | -27.2 ± 3.89 *P* < 0.001 | -54.7 ± 5.73 *P* < 0.001 | -24.2 ± 5.05 *P* = 0.001 | -5.70 ± 1.899 *P* = 0.013 |  | 19, 21 | 78.9 | 321 | 329 | 0.787 | 19.3 |

The covariates follow the same order as they are presented in the models.

Type = cattle type (dairy or beef). Cattle type was included as 2 binary category in the models: Dairy was coded as 1 if the animal was dairy cattle and as 0 if otherwise, and Beef was coded as 1 if the animal was beef cattle and as 0 if otherwise.

All variables (except cattle type) were centered to their mean value within each cattle type presented in Table 1 in the main document.

Outliers ID = Individual identification of outlier data points using the CCC-based method. Studies references and their IDs are listed in Figure 2 in the main document.

Bromoform (CHBr_3_) dose (mg/kg DM) was natural log-transformed before models fitting.

*I*^2^ = proportion of total variability due to heterogeneity.

AIC, BIC = Akaike and Bayesian information criteria.

CCC = Concordance Correlation Coefficient.

RMSE = root mean square error using leave-one-out cross-validation, quantifying how far the model's predictions typically deviate from the actual observed values, in percentage points of relative mean difference (RMD).

### **Table S6.** Comparison of regression models evaluating the effect of bromoform (CHBr_3_; in LN mg CHBr_3_/kg DM) dose and additional dietary or animal covariates on the relative mean difference in CH_4_ yield (estimates ± SE) with identification of outliers using the CCC-based method

|  |  | Estimates | | | |  | Model metrics | | | | | |
| --- | --- | --- | --- | --- | --- | --- | --- | --- | --- | --- | --- | --- |
| Models |  | Dairy | Beef | 2nd  Covariate | 3rd  Covariate |  | Outliers  ID | *I*^2^ | AIC | BIC | CCC | RMSE (%) |
| Type + CHBr_3_ |  | -23.2 ± 2.29 *P* < 0.001 | -50.6 ± 6.04 *P* < 0.001 | -20.5 ± 2.91 *P* < 0.001 | NA |  | 13, 24 | 80.9 | 313 | 319 | 0.732 | 18.2 |
| Type + CHBr_3_ + (CHBr_3_)^2^ |  | -23.2 ± 2.42 *P* < 0.001 | -50.5 ± 6.94 *P* < 0.001 | -20.6 ± 4.81 *P* = 0.002 | -0.217 ± 2.7444 *P* = 0.939 |  | 13, 24 | 81.0 | 315 | 323 | 0.732 | 21.0 |
| Type + CHBr_3_ + DMI |  | -23.0 ± 2.33 *P* < 0.001 | -50.5 ± 6.32 *P* < 0.001 | -20.6 ± 2.88 *P* < 0.001 | -2.57 ± 1.996 *P* = 0.230 |  | 13, 24 | 79.3 | 312 | 320 | 0.755 | 18.1 |
| Type + CHBr_3_ + CP |  | -23.2 ± 2.42 *P* < 0.001 | -44.4 ± 3.46 *P* < 0.001 | -19.1 ± 3.06 *P* < 0.001 | 0.237 ± 1.3743 *P* = 0.867 |  | 13, 14, 19, 21, 24 | 63.3 | 267 | 275 | 0.800 | 13.9 |
| Type + CHBr_3_ + NDF |  | -23.3 ± 2.28 *P* < 0.001 | -46.7 ± 3.84 *P* < 0.001 | -19.2 ± 2.47 *P* < 0.001 | 0.444 ± 0.2172 *P* = 0.071 |  | 13, 19,  21, 24 | 61.8 | 279 | 286 | 0.796 | 14.9 |
| Type + CHBr_3_ + Starch |  | -22.5 ± 2.49 *P* < 0.001 | -56.0 ± 5.46 *P* < 0.001 | -23.4 ± 3.74 *P* < 0.001 | -0.798 ± 0.2258 *P* = 0.010 |  | 10, 11, 24 | 71.6 | 275 | 283 | 0.868 | 15.2 |
| Type + CHBr_3_ + FAT |  | -23.1 ± 3.32 *P* < 0.001 | -50.7 ± 6.19 *P* < 0.001 | -21.5 ± 4.89 *P* = 0.002 | -4.91 ± 1.517 *P* = 0.010 |  | 19, 21 | 81.3 | 314 | 322 | 0.757 | 19.0 |

The covariates follow the same order as they are presented in the models.

Type = cattle type (dairy or beef). Cattle type was included as 2 binary category in the models: Dairy was coded as 1 if the animal was dairy cattle and as 0 if otherwise, and Beef was coded as 1 if the animal was beef cattle and as 0 if otherwise.

All variables (except cattle type) were centered to their mean value within each cattle type presented in Table 1 in the main document.

Outliers ID = Individual identification of outlier data points using the CCC-based method. Studies references and their IDs are listed in Figure 2 in the main document.

Bromoform (CHBr_3_) dose (mg/kg DM) was natural log-transformed before models fitting.

*I*^2^ = proportion of total variability due to heterogeneity.

AIC, BIC = Akaike and Bayesian information criteria.

CCC = Concordance Correlation Coefficient.

RMSE = root mean square error using leave-one-out cross-validation, quantifying how far the model's predictions typically deviate from the actual observed values, in percentage points of relative mean difference (RMD).

### **Table S7.** Comparison of regression models evaluating the effect of bromoform (CHBr_3_; in LN mg CHBr_3_/kg DM) dose and additional dietary or animal covariates on the relative mean difference in CH_4_ intensity (estimates ± SE) with identification of outliers using the CCC-based method

|  |  | Estimates | | | |  | Model metrics | | | | | |
| --- | --- | --- | --- | --- | --- | --- | --- | --- | --- | --- | --- | --- |
| Models |  | Dairy | Beef | 2nd  Covariate | 3rd  Covariate |  | Outliers  ID | *I*^2^ | AIC | BIC | CCC | RMSE (%) |
| Type + CHBr_3_ |  | -26.2 ± 2.98 *P* < 0.001 | -52.1 ± 7.09 *P* < 0.001 | -22.2 ± 6.24 *P* = 0.009 | NA |  | 19, 20 | 59.2 | 235 | 241 | 0.761 | 18.1 |
| Type + CHBr_3_ + (CHBr_3_)^2^ |  | -25.4 ± 3.15 *P* < 0.001 | -48.8 ± 7.62 *P* = 0.001 | -25.6 ± 5.98 *P* = 0.005 | -5.08 ± 2.493 *P* = 0.088 |  | 19, 20 | 59.6 | 235 | 242 | 0.795 | 20.6 |
| Type + CHBr_3_ + DMI |  | -26.1 ± 3.77 *P* < 0.001 | -48.4 ± 5.53 *P* < 0.001 | -23.9 ± 7.24 *P* = 0.016 | -0.715 ± 2.7373 *P* = 0.803 |  | 19, 20, 21 | 46.3 | 221 | 227 | 0.788 | 18.8 |
| Type + CHBr_3_ + CP |  | -26.1 ± 3.37 *P* < 0.001 | -51.7 ± 6.91 *P* < 0.001 | -22.0 ± 6.57 *P* = 0.016 | -0.662 ± 2.7065 *P* = 0.815 |  | 19, 20 | 60.8 | 237 | 244 | 0.772 | 18.5 |
| Type + CHBr_3_ + NDF |  | -27.3 ± 2.54 *P* < 0.001 | -45.4 ± 5.00 *P* < 0.001 | -22.8 ± 3.76 *P* = 0.001 | 0.960 ± 0.4331 *P* = 0.069 |  | 19, 20, 21 | 17.2 | 209 | 216 | 0.844 | 14.1 |
| Type + CHBr_3_ + Starch |  | -25.5 ± 1.99 *P* < 0.001 | -46.5 ± 2.70 *P* < 0.001 | -24.0 ± 4.48 *P* = 0.003 | -0.579 ± 0.1689 *P* = 0.019 |  | 19, 20, 21 | 11.9 | 191 | 197 | 0.867 | 13.3 |
| Type + CHBr_3_ + FAT |  | -26.4 ± 2.80 *P* < 0.001 | -49.8 ± 8.49 *P* = 0.001 | -21.5 ± 6.17 *P* = 0.013 | -4.80 ± 3.303 *P* = 0.196 |  | 19, 20 | 50.4 | 234 | 240 | 0.786 | 17.7 |

The covariates follow the same order as they are presented in the models.

Type = cattle type (dairy or beef). Cattle type was included as 2 binary category in the models: Dairy was coded as 1 if the animal was dairy cattle and as 0 if otherwise, and Beef was coded as 1 if the animal was beef cattle and as 0 if otherwise.

All variables (except cattle type) were centered to their mean value within each cattle type presented in Table 1 in the main document.

Outliers ID = Individual identification of outlier data points using the CCC-based method. Studies references and their IDs are listed in Figure 2 in the main document.

Bromoform (CHBr_3_) dose (mg/kg DM) was natural log-transformed before models fitting.

*I*^2^ = proportion of total variability due to heterogeneity.

AIC, BIC = Akaike and Bayesian information criteria.

CCC = Concordance Correlation Coefficient.

RMSE = root mean square error using leave-one-out cross-validation, quantifying how far the model's predictions typically deviate from the actual observed values, in percentage points of relative mean difference (RMD).

### **Table S8.** Comparison of regression models evaluating the effect of bromoform (CHBr_3_) dose on the relative mean difference in DMI and products (milk, ECM, ADG) (estimates ± SE) with identification of outliers using the CCC-based method

|  |  | Estimates | | |  | Model metrics | | | | | |
| --- | --- | --- | --- | --- | --- | --- | --- | --- | --- | --- | --- |
| Models |  | Dairy | Beef | CHBr_3_ |  | Outliers ID | *I*^2^ | AIC | BIC | CCC | RMSE (%) |
| *DMI* |  |  |  |  |  |  |  |  |  |  |  |
| Type + CHBr_3_ |  | -6.45 ± 1.588 *P* = 0.002 | -3.26 ± 1.169 *P* = 0.018 | -0.269 ± 0.0410 *P* < 0.001 |  | 20, 31 | 39.5 | 232 | 238 | 0.643 | 5.73 |
| *Products* |  |  |  |  |  |  |  |  |  |  |  |
| Type + CHBr_3_ |  | -4.66 ± 2.571 *P* = 0.103 | 3.70 ± 2.784 *P* = 0.217 | -0.271 ± 0.0724 *P* = 0.005 |  | 18, 20, 25, 26 | 58.8 | 238 | 244 | 0.280 | 10.2 |

The covariates follow the same order as they are presented in the models. The first covariate corresponds to the cattle type which must be one of the two columns: Dairy or Beef.

Type = cattle type (dairy or beef). Cattle type was included as 2 binary category in the models: Dairy was coded as 1 if the animal was dairy cattle and as 0 if otherwise, and Beef was coded as 1 if the animal was beef cattle and as 0 if otherwise.

Bromoform (CHBr_3_) dose (mg/kg DM) was centered to their mean value within cattle type presented in Table 2 in the main document.

Outliers ID = Individual identification of outlier data points using the CCC-based method. Studies references and their IDs are listed in Figure 2 in the main document.

*I*^2^ = proportion of total variability due to heterogeneity.

AIC, BIC = Akaike and Bayesian information criteria.

CCC = Concordance Correlation Coefficient.

RMSE = root mean square error using leave-one-out cross-validation, quantifying how far the model's predictions typically deviate from the actual observed values, in percentage points of relative mean difference (RMD).

**Example: Calculating change (%) in CH_4_ production**

**Scenario:**

We want to calculate the percentage change in CH_4_ production for a dairy cow with the following characteristics:

CHBr_3_ dose: 20 mg/kg DM

Dietary starch content: 15% DM

**Equation:**

$$\mathrm{Change}\left( \% \right)\mathrm{in}\mathrm{CH}_{4}\mathrm{production}=(Dairy or Beef estimate)-26.6 \pm3.75\times(LN\left( \mathrm{CHBr}_{3} dose)-mean LN(\mathrm{CHBr}_{3} dose) \right)-0.764 \pm0.1525\times(Starch-\mathrm{mean}Starch),$$

**Step 1: Gather input values**

Dairy estimate: −24.8 (from Table 3 in the main document)

Mean LN(CHBr_3_): 2.685 (from Table 1 in the main document)

Mean starch: 12.8% DM (from Table 1 in the main document)

**Step 2: Calculate LN of CHBr_3_ dose**

LN(CHBr_3_) = LN(20) = 2.996 (Important: Use at least three decimal places)

**Step 3: Compute each term in the equation**

CHBr_3_ term: −26.6 × (LN(CHBr_3_) − mean LN(CHBr_3_))

= −26.6 × (2.996 − 2.685)
= −26.6 × 0.311
= −8.27

Starch term: −0.764 × (Starch − mean Starch)
= −0.764 × (15 − 12.8)
= −0.764 × 2.2
= −1.68

**Step 4: Sum all terms**

Change (%) in CH₄ production = −24.8 (Dairy estimate) – 8.27 (CHBr_3_ term) − 1.68 (Starch term)
= −24.8 – 8.27 − 1.68
= −34.8

**Interpretation:**

The model predicts a 34.8% reduction in CH_4_ production for a dairy cow with:

CHBr_3_ dose = 20 mg/kg DM

Dietary starch = 15% DM
